# Optimised DNA isolation from marine sponges for natural sampler DNA (nsDNA) metabarcoding

**DOI:** 10.1101/2022.07.11.499619

**Authors:** Lynsey R. Harper, Erika F. Neave, Graham S. Sellers, Alice V. Cunnington, María Belén Arias, Jamie Craggs, Barry MacDonald, Ana Riesgo, Stefano Mariani

**Author notes:** Corresponding author: Stefano Mariani.

## Abstract

Marine sponges have recently been recognised as natural samplers of environmental DNA (eDNA) due to their effective water filtration and their ubiquitous, sessile and regenerative nature. However, laboratory workflows for metabarcoding of sponge tissue have not been optimised to ensure that these natural samplers achieve their full potential for community survey. We used a phased approach to investigate the influence of DNA isolation procedures on the biodiversity information recovered from sponges. In Phase 1, we compared three treatments of residual ethanol preservative in sponge tissue alongside five DNA extraction protocols. The results of Phase 1 informed which ethanol treatment and DNA extraction protocol should be used in Phase 2, where we assessed the effect of starting tissue mass on extraction success and whether homogenisation of sponge tissue is required. Phase 1 results indicated that ethanol preservative may contain unique and/or additional biodiversity information to that present in sponge tissue, but blotting tissue dry generally recovered more taxa and generated more sequence reads from the wild sponge species. Tissue extraction protocols performed best in terms of DNA concentration, taxon richness and proportional read counts, but the non-commercial tissue protocol was selected for Phase 2 due to cost-efficiency and greater recovery of target taxa. In Phase 2 overall, we found that homogenisation may not be required for sponge tissue and more starting material does not necessarily improve taxon detection. These results combined provide an optimised DNA isolation procedure for sponges to enhance marine biodiversity assessment using natural sampler DNA metabarcoding.

## 1. Introduction

The emergence of environmental DNA (eDNA) analysis signalled a new age of biodiversity monitoring, facilitating faster, non-invasive, cost-efficient and comprehensive surveys for single species and whole communities than was previously thought possible (Lawson Handley 2015; Deiner et al., 2017; Miya 2021). Such has been the rapid, exponential growth of this field in the last decade (Tsuji et al., 2019; Thalinger et al., 2021) that it is now beginning to branch. In addition to the now relatively standard analyses of aquatic eDNA and soil/sediment eDNA, studies have been published on species detection using eDNA extracted from snow (Franklin et al., 2019; Kinoshita et al., 2019) and air (Clare et al., 2021; Lynggaard et al., 2022; Roger et al., 2022). Innovative research has also demonstrated that water can be used to aggregate eDNA deposited on plants (Valentin et al., 2020) or fishing nets (Russo et al., 2020) for species detection. Less closely intertwined but stemming from the same roots is natural sampler DNA (nsDNA) (Siegenthaler et al., 2019), which refers to the analysis of DNA collected by live organisms. This is not a novel concept as researchers have been analysing the DNA present in faeces and gut contents of consumers (Pompanon et al., 2012; Nørgaard et al., 2021) and hematophagous insects (Schnell et al., 2012; Massey et al., 2021) for years, but more creative applications are now being explored, including flowers and cow dung as samplers of insect assemblages (Thomsen & Sigsgaard, 2019; Sigsgaard et al., 2020; Harper et al., 2022), dung beetles as samplers of mammal communities (Drinkwater et al., 2021), and sponges as samplers of marine biodiversity (Mariani et al., 2019; Turon et al., 2020; Jeunen et al., 2021).

Marine sponges can filter thousands of litres of water in a single day (Gökalp et al., 2020), vastly exceeding the volume that could be collected with artificial devices for aquatic eDNA metabarcoding. As water is filtered, particles are trapped and concentrated in sponge tissue until digestion or excretion, meaning that a biopsy can be taken to provide DNA for metabarcoding analysis (Mariani et al., 2019; Turon et al., 2020; Jeunen et al., 2021). Sponge nsDNA metabarcoding offers several advantages over aquatic eDNA metabarcoding. The high-tech robotics being used to sample large volumes of water for aquatic eDNA (e.g. automated underwater vehicles) are expensive to build and operate, are ineffective for inaccessible and/or complex habitats, and often target a limited set of taxa (Mariani et al., 2019). Although passive filtration has been explored as an alternative to active filtration using peristaltic or vacuum pumps, they require long submergence times and have saturation limits (Kirtane et al., 2020; Bessey et al., 2021a, b). Sponges are present in nearly every aquatic habitat, and sample collection is inexpensive and non-destructive with minimal environmental impact. Provided that biopsies are not destructive and conducted carefully, sponges will regenerate quickly. Therefore, sponges offer a low-tech, affordable and standardised method of surveying marine biodiversity, and could serve as long-term monitoring stations (Mariani et al., 2019). However, there is much work to be done before sponges can be reliably and routinely used in community surveys, specifically sampling and laboratory protocols (Jeunen et al., 2021).

Two sponge natural sampler DNA studies to date (Mariani et al., 2019; Turon et al., 2020) utilised samples from existing collections and extracted DNA from tissue with commercial kits. The other study (Jeunen et al., 2021) compared dive surveys and aquatic eDNA to sponge tissue and the ethanol used to preserve sponge tissue, and also extracted DNA using a commercial kit. It remains undetermined whether certain sponge species are better samplers than others and if this is related to phenotype. The spatial extent of biodiversity information recovered by a single sponge is yet to be ascertained along with the optimal mass/volume for biopsies to maximise data generation without being lethal to the sponge sampled (Mariani et al., 2019). The influence of DNA extraction method on biodiversity detection from sponge tissues is unknown (Jeunen et al., 2021), but this has been found to affect detection success with aquatic eDNA (Deiner et al., 2018; Jeunen et al., 2019), bulk invertebrates (Hermans et al., 2018; Majaneva et al., 2018) and faecal samples (Kaunisto et al., 2017). Furthermore, it has not been investigated if samples should be homogenised to ensure DNA is evenly distributed throughout the sample prior to extraction, similar to bulk invertebrate and faecal samples (Gosselin et al., 2017; Pereira-da-Conceicoa et al., 2020). The optimal amount of starting material to maximise species recovery without introducing PCR inhibitors is also unknown, but using larger amounts of soil was found to improve invertebrate detection (Kirse et al., 2021).

Here, we focus on the influence of laboratory workflows on the performance of sponge natural sampler DNA metabarcoding, specifically DNA isolation procedures. We used a phased approach with three sponge species to investigate whether residual ethanol preservative, DNA extraction protocol, type of starting material, and amount of starting material influence sponge nsDNA metabarcoding performance using the following extraction success criteria: 1) total DNA yield, 2) target DNA concentration, 3) taxon richness, 4) proportional read counts, and 5) community composition. In Phase 1, we compared three treatments of residual ethanol preservative in sponge tissue (no removal, removal via blotting, removal via centrifugation) alongside five DNA extraction protocols, including commercial kits and non-commercial protocols. Using the optimal ethanol treatment and DNA extraction protocol from Phase 1, we compared different volumes of homogenised sponge tissue to different weights of dried sponge tissue. We synthesise the results from each experimental phase to provide an optimised DNA isolation procedure for sponge tissue to enhance marine biodiversity assessment using natural sampler DNA metabarcoding.

## 2. Materials and Methods

### 2.1 Workspace and decontamination procedures

A unidirectional workflow was used for sample processing. Sponge tissue was stored and handled in a laboratory dedicated to the processing of environmental samples with low DNA concentrations. Prepared PCR reactions were transported to separate laboratories for: addition of PCR positive controls and PCR amplification; gel electrophoresis, library preparation, and storage; and library quality checks, quantification, and sequencing. Bench space and non-immersible equipment in all laboratories were sterilised before and after use by wiping with fresh paper towel and 10% v/v bleach solution (made using Cleanline thin bleach containing 4.53% sodium hypochlorite), followed by 70% v/v ethanol solution. Immersible equipment was sterilised in a 10% v/v bleach bath for at least 10 minutes, then immersed in a 5% v/v Lipsol detergent bath and rinsed with deionised water. Bleach and Lipsol detergent baths were changed weekly or daily during periods of heavy use. Stainless steel 5 mm beads (Qiagen, Germany) were additionally sterilised by heating at 220°C for 3 hours. Plastics and hoods were sterilised with UV light for at least 30 minutes. Full details of decontamination procedures are provided in Supporting Information.

### 2.2 Sponge selection

For this experiment, we used samples from three marine sponges which are the focus of ongoing research employing nsDNA metabarcoding: 1) an invasive sponge, *Lendenfeldia chondrodes* (Galitz et al., 2018), present in the aquarium facility of the Horniman Museum and Gardens, London, 2) *Vazella pourtalesii*, which is a siliceous sponge that colonises hydroacoustic moorings in Nova Scotia, Canada, and 3) *Phakellia ventilabrum*, which is a Bubarid sponge that forms relatively dense aggregations on rock-sand habitats and is broadly distributed across the North Atlantic. *L. chondrodes* provided an opportunity to groundtruth natural sampler DNA metabarcoding in a controlled tank environment with 25 species of known abundance, whereas *V. pourtalesii* and *P. ventilabrum* were collected from natural marine environments with more diverse species assemblages and differed in phenotype as well as filtration efficiency. *L. chondrodes* and *V. pourtalesii* were used in Phase 1, whereas *V. pourtalesii* and *P. ventilabrum* were used in Phase 2 due to the greater amounts of starting material required for DNA extraction.

### 2.3 Sponge collection

All sponges were detached from their substrate using a sterile diving knife or sterile disposable surgical scalpel (Swann-Morton No. 21, Fisher Scientific, UK) whilst wearing disposable gloves. Individuals of *L. chondrodes* were small thus an ~1 cm^3^ biopsy was taken. Each biopsy was placed in a 2 ml DNA LoBind tube (Eppendorf, Fisher Scientific, UK) containing 1 ml of molecular grade 100% ethanol (Fisher Scientific, UK) and stored at −20°C for transport to Liverpool John Moores University (LJMU). At LJMU, forceps were used to transfer each biopsy to a new 2 ml DNA LoBind tube containing 1 ml of molecular grade 100% ethanol as water present in tissue may have diluted the original ethanol. Samples were stored at −20°C for three months until DNA extraction. As *V. pourtalesii* colonises hydroacoustic moorings, samples were collected during mooring retrieval undertaken by Ocean Tracking Network and Fisheries and Oceans Canada in July 2021. Whole individuals were removed from the moorings, placed in sterile pots containing 500 ml of molecular grade 100% ethanol, and stored at −20°C once research vessels returned to shore. The ethanol in each pot was changed twice before samples were shipped to LJMU and stored at −20°C for three months until DNA extraction. For *P. ventilabrum,* a ~5 cm^3^ biopsy was taken from each individual sampled from Rockall Bank (coordinates: 57.1705, −13.638) at 185-189 m depth in May 2018, then placed in a 50 ml falcon tube containing 25 ml of 100% ethanol and stored at 4°C. Again, the ethanol was changed twice before samples were shipped to LJMU in January 2021 where they were stored at −20°C for three months until DNA extraction.

### 2.4 DNA extraction

#### 2.4.1 Phase 1

Three *L. chondrodes* samples and three *V. pourtalesii* samples (each from different moorings) were used in Phase 1 (Fig. 1). Each sample from each sponge was subjected to a different ethanol treatment as ethanol is known to interact with chemicals used during DNA extraction and inhibit PCR amplification, but residual ethanol preservative may be present in porous sponge tissue. The ethanol treatments were: 1) no removal of residual ethanol (wet treatment), 2) removal of residual ethanol by blotting sponge tissue against filter paper inside a petri dish (blot treatment), and 3) biopsies were transferred to a new 2 ml DNA LoBind tube and centrifuged for 1 minute at maximum speed, following which the ethanol was removed with a pipette and centrifugation repeated until no more ethanol was observed (spin treatment). After each ethanol treatment, samples were cut into small pieces within a petri dish using dissection scissors. Pieces were added to a weigh boat until a maximum weight of 500 mg was observed. Each subsample was transferred to a high-impact 2 ml screw cap microtube (Starlab, UK) containing a 5 mm stainless steel bead and twice the amount of 1x TE buffer to tissue weight was added to each microtube (e.g. 500 μl to 250 mg). This resulted in 20-30 subsamples per *V. pourtalesii* sample. Each *L. chondrodes* sample weighed ~500 mg individually thus no subsamples were required. The (sub)samples were homogenised on a Qiagen TissueLyser II for 2 minutes at maximum speed. Using a 1000 μl pipette tip, the homogenate from each subsample was pooled in a 15 ml falcon tube corresponding to each original sample. Twelve 200 μl aliquots of each *V. pourtalesii* sample and twelve 50 μl aliquots of each *L. chondrodes* sample (Fig. 1) were transferred to individual 2 ml DNA LoBind tubes and frozen at −20°C alongside the homogenised samples.

**Figure 1.**
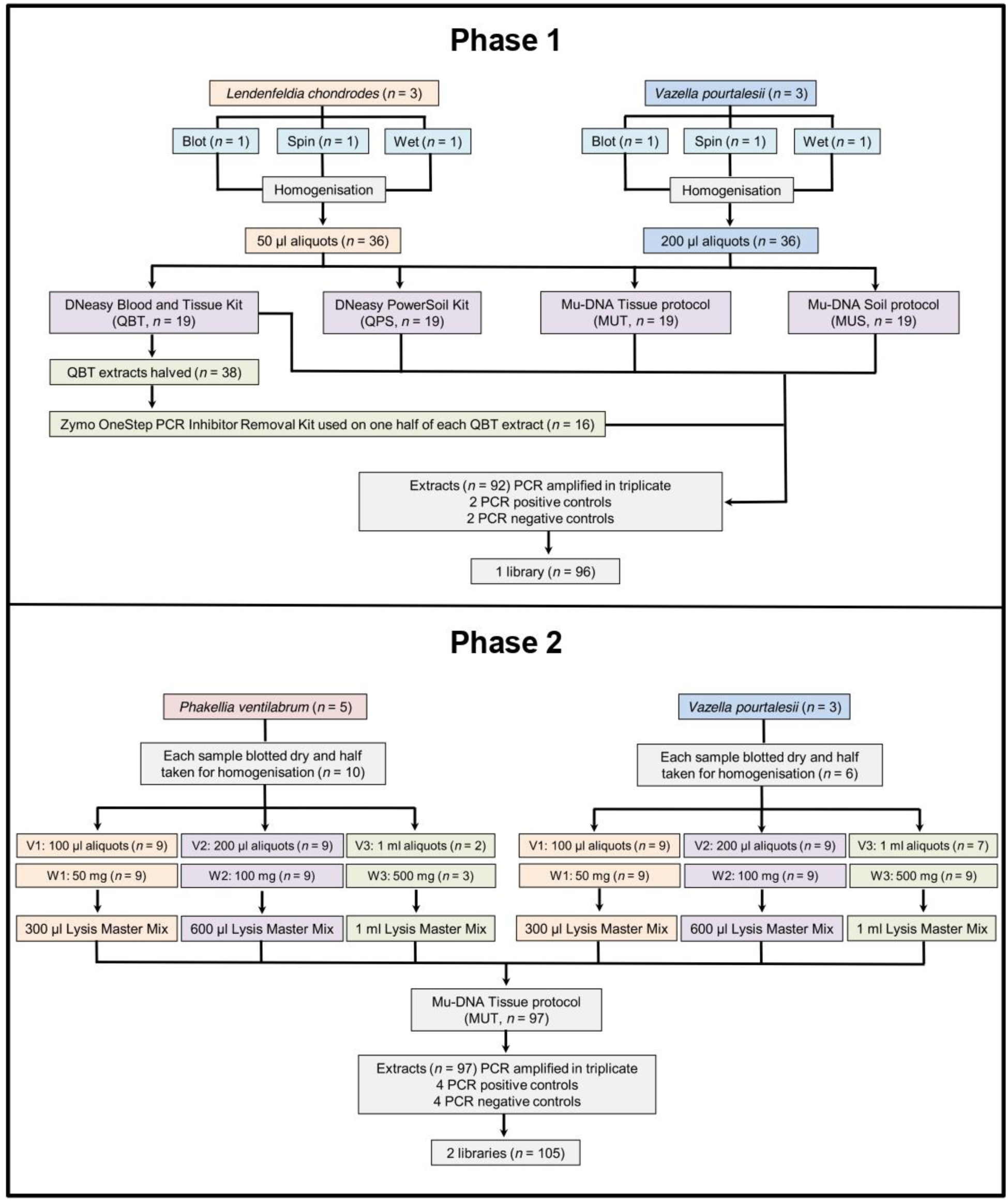
Flow diagram illustrating the laboratory workflow for the phased experiment.

The aliquots were used as the starting material for extraction with three commercial and two non-commercial DNA extraction protocols: 1) Qiagen DNeasy PowerSoil Kit (QPS), 2) Qiagen DNeasy Blood and Tissue Kit (QBT), 3) Qiagen DNeasy Blood and Tissue Kit and post-extraction inhibitor removal with the Zymo Research OneStep™ PCR Inhibitor Removal Kit (QBT-ZYMO), 4) Mu-DNA soil protocol (MUS), and 5) Mu-DNA tissue protocol (MUT) with an added inhibitor removal step (Sellers et al., 2018) (Fig. 1). The QPS, QBT, MUS and MUT protocols were used on three aliquots of each sample (*n* = 72) following the manufacturer’s instructions or steps outlined in Sellers et al. (2018) with one modification. Elution buffer (100 μl) was added to each spin column, incubated for 5 minutes at room temperature before centrifugation for 1 minute at 10,000 x g, then the eluate was collected and added to the same spin column again for incubation for 5 minutes at room temperature before centrifugation for 1 minute at 10,000 x g. An extraction blank (extraction buffers only) was processed alongside samples for each extraction protocol (*n* = 4).

The extracts resulting from QBT extraction were halved (Fig. 1), and one half was processed with the Zymo Research OneStep™ PCR Inhibitor Removal Kit (Cambridge Bioscience, UK) for the QBT-ZYMO protocol (*n* = 19). Notably, three preps from the OneStep™ PCR Inhibitor Removal Kit failed, resulting in only 16 extracts for the QBT-ZYMO protocol (Fig. 1). A 25 μl and 5 μl aliquot of each extract (*n* = 92) were transferred to PCR strip tubes (Starlab, UK). All original DNA extracts and 25 μl aliquots were frozen at −20°C until PCR preparation. A Nanodrop was used to measure DNA purity and a Qubit™ 4 Fluorometer (Thermo Scientific, UK) with Qubit™ dsDNA HS Assay Kit (Fisher Scientific, UK) was used to measure total DNA yield from the 5 μl aliquots, which were subsequently discarded.

#### 2.4.2 Phase 2

Three *V. pourtalesii* samples (each from different moorings) and five *P. ventilabrum* samples (four from one location and one from another location) were used in Phase 2 (Fig. 1). Based on the findings of Phase 1 (see Results), all samples were blotted against filter paper in a petri dish to remove residual ethanol preservative and extracted with the MUT protocol. Samples from each sponge species were halved (*n* = 12), with each half being used to provide either dry tissue or homogenate as starting material (Fig. 1). Homogenisation of each sample was performed as described in Phase 1. Three dry tissue weights and three homogenate volumes were compared in triplicate for each sample (*n* = 108), and the volume of Lysis Master Mix (1 ml contains 730 μl of Lysis Solution, 250 μl of Tissue Lysis Additive and 20 μl of Proteinase K) for the MUT protocol was adjusted proportionately: 1) 50 mg and 100 μl (≈50 mg) to 300 μl of Lysis Master Mix, 2) 100 mg and 200 μl (≈100 mg) to 600 μl of Lysis Master Mix, and 3) 500 mg and 1000 μl (≈500 mg) to 1 ml of Lysis Master Mix. Extraction followed the steps outlined in Sellers et al. (2018) with the same modification for elution as Phase 1. Notably, there was not enough tissue for some samples to obtain triplicate 500 mg weights or 1000 μl volumes, even when samples collected from the same location were used as replicates (PHAK4, PHAK5), resulting in 105 DNA extracts. An extraction blank (extraction buffers only) was included for each round of extractions (*n* = 4) (Fig. 1). A 25 μl and 5 μl aliquot of each extract (*n* = 109) were transferred to PCR strip tubes (Starlab, UK). All original DNA extracts and 25 μl aliquots were frozen at −20°C until PCR preparation. A Nanodrop was used to measure DNA purity and a Qubit™ 4 Fluorometer was used to measure total DNA yield.

### 2.5 PCR amplification and library preparation

DNA extracts were PCR amplified using the Tele02 primers (F: 5’-AAACTCGTGCCAGCCACC-3’, R: 5’-GGGTATCTAATCCCAGTTTG-3’) which target a ~167 bp fragment of the mitochondrial 12S ribosomal RNA (rRNA) gene and achieved >98% teleost species detection when tested *in silico* (Taberlet *et al.,* 2018). Although the primers were designed to target teleost fish, mammal and bird DNA may also amplify (Mariani *et al.,* 2021). To ease sample demultiplexing and mitigate cross-contamination and/or tag switching during sequencing, each sample was PCR amplified using the locus primers attached to a unique 8 bp tag shared by the forward and reverse primer. Each tag (*n* = 96) differed by at least three base pairs from other tags and included 2-4 degenerate bases (Ns) at the beginning of the tag sequence to improve clustering during initial sequencing. For Phase 2, samples were processed in two batches to facilitate use of the same tags in different pools for adapter ligation.

Each sample was PCR amplified in triplicate using 20 μl reactions consisting of 10 μl of MyFi™ Mix (Meridian Bioscience, UK), 1 μl of each forward and reverse primer (10 μM, Eurofins), 0.16 μl of Bovine Serum Albumin (20 mg/ml, Thermo Scientific, UK), 5.84 μl of UltraPure™ Distilled Water (Invitrogen, UK), and 2 μl of DNA template. Two PCR positive controls (iridescent catfish, *Pangasianodon hypophthalmus*, tissue DNA at 0.05 ng/μl) and two PCR negative controls (molecular grade water) were included in each PCR run (up to 96 reactions). PCR was performed on a T100 Thermal Cycler (Bio-Rad Laboratories Ltd, UK) with the following profile: 95°C for 10 minutes, followed by 40 cycles of 95°C for 30 s, 60°C for 45 s, and 72°C for 30 s, and final elongation at 72°C for 5 minutes. PCR products were run on a 2% agarose gel, made with 150 ml of 1x TBE buffer (Invitrogen, UK), 3 g of agarose powder (Fisher Scientific, UK), and 1.5 μl of SYBR™ Safe DNA Gel Stain (Invitrogen, UK), at 100 V for 40 minutes using 1 μl of PCR product to 1 μl of 10x BlueJuiceTM Gel Loading Buffer (Invitrogen, UK). Gels were visualised with a Bio-Rad Gel Doc™ EZ Imaging System.

PCR replicates for each sample were pooled prior to purification with Mag-Bind^®^ Total Pure NGS Beads (Omega Bio-tek Inc., GA, USA), following the double size selection protocol established by Bronner et al. (2009). Ratios of 1x and 0.6x magnetic beads to 30 μl of PCR product were used. Eluted DNA (20 μl) was stored at −20°C until quantification using a Qubit™ 4 Fluorometer. Purified samples were normalised and pooled in equimolar concentration. Phase 1 consisted of one pool of 96 samples/controls. Phase 2 required two pools - the first consisted of 65 samples/controls from Phase 2 and the second consisted of 40 samples/controls from Phase 2 and 25 samples/controls from other projects. Sample pools were purified using the aforementioned bead ratios and elution in 25 μl, then concentrated using a 1x bead ratio and elution in 45 μl.

End repair, adapter ligation and PCR were performed using the NEXTFLEX^®^ Rapid DNA-Seq Kit 2.0 for Illumina^®^ Platforms (PerkinElmer, UK) according to the manufacturer’s protocol. An Agilent 2200 TapeStation and High Sensitivity D1000 ScreenTape (Agilent Technologies, UK) indicated secondary product remained, thus gel extraction was performed on each pool using the GeneJET Gel Extraction Kit (Thermo Scientific, UK) with elution in 20 μl. Each pool was quantified using quantitative PCR (qPCR) on a Rotor-Gene Q (Qiagen, Germany) with the NEBNext^®^ Library Quant Kit for Illumina^®^ (New England Biolabs, UK) and diluted to 1 nM. For Phase 2, 6 μl of each pool were combined into one library. The final libraries and PhiX Control were quantified using qPCR before the Phase 1 library was sequenced at 50 pM with 10% PhiX Control and the Phase 2 library was sequenced at 60 pM with 10% PhiX Control on an Illumina^®^ iSeq™ 100 using iSeq™ 100 i1 Reagent v2 (300-cycle) (Illumina Inc., CA, USA).

### 2.6 Bioinformatics

#### 2.6.1 Reference databases

Sequence data were processed in two batches to facilitate taxonomic assignment against different reference databases. Sequences for *L. chondrodes* samples and associated controls were compared against a reference database for the aquarium fish species, the PCR positive control, and common domestic species (human *Homo sapiens*, cow *Bos taurus*, pig *Sus scrofa*, chicken *Gallus gallus*, sheep *Ovis aries*, dog *Canis lupus*, and cat *Felis catus*). Sequences for *V. pourtalesii* and *P. ventilabrum* samples and associated controls were compared against a marine vertebrate reference database, the PCR positive control, and common domestic species. Full details of database creation can be found in the Supporting Information.

#### 2.6.2 Data processing and taxonomic assignment

Sequence data were automatically demultiplexed to separate (forward and reverse) fastq files per library using the onboard Illumina MiSeq Reporter software. Library sequence reads were further demultiplexed to sample using a custom Python script. *Tapirs*, a reproducible workflow for the analysis of DNA metabarcoding data (https://github.com/EvoHull/Tapirs), was used for processing and taxonomic assignment of demultiplexed sequence reads. *Tapirs* uses the *Snakemake* workflow manager (Köster & Rahmann, 2012) and a *conda* virtual environment to ensure software compatibility.

Raw reads were quality trimmed from the tail with a 5 bp sliding window (qualifying phred score of Q30 and an average window phred score of Q30) using fastp (Chen et al., 2018), allowing no more than 40% of the final trimmed read bases to be below Q30. Poly X/G tail trimming (≥ 5 bp) removed any remaining Illumina iSeq™ 100 sequencing artefacts. Primers were removed by trimming the first 18 bp and 20 bp of the forward and reverse reads respectively. Reads were then tail cropped to a maximum length of 150 bp and reads shorter than 90 bp were discarded. Sequence read pairs were merged into single reads using fastp, provided there was a minimum overlap of 20 bp, no more than 5% mismatches and no more than 5 mismatched bases between pairs. Only forward reads were kept from read pairs that failed to be merged. A final length filter removed any reads longer than 200 bp to ensure sequence lengths approximated the expected fragment size (~167 bp). Redundant sequences were removed by clustering at 100% read identity and length (--derep_fulllength) in VSEARCH (Rognes et al., 2016). Clusters represented by less than three sequences were omitted from further processing. Reads were further clustered (--cluster_unoise) to remove redundancies due to sequencing errors (retaining all cluster sizes). Retained sequences were screened for chimeric sequences with VSEARCH (--uchime3_denovo). The final clustered, non-redundant query sequences were then compared against the corresponding reference database using *BLAST* (Zhang et al., 2000). Taxonomic identity was assigned using a custom majority lowest common ancestor (MLCA) approach based on the top 2% query *BLAST* hit bit-scores, with at least 90% query coverage. A minimum *BLAST* hit identity filter of 80% was used for *L. chondrodes* samples and the aquarium fish reference database, whereas a minimum identity of 98% was used for *V. pourtalesii* and *P. ventilabrum* samples and the marine vertebrate reference database. Of these filtered hits for both batches, 80% of unique taxonomic lineages therein had to agree at descending taxonomic rank (domain, phylum, class, order, family, genus, species) for it to be assigned a taxonomic identity. If a query had a single *BLAST* hit, it was assigned directly to this taxon only if it met all previous MLCA criteria. Read counts assigned to each taxonomic identity were calculated from query cluster sizes. Lowest taxonomic rank was to species and assignments higher than order were classed as unassigned.

### 2.7 Data analysis

#### 2.7.1 Dataset refinement

Taxonomically assigned data was combined for downstream processing in R v3.6.3 (R Core Team, 2020). All plots were produced using the packages ggplot2 v3.3.6 (Wickham, 2016) and ggpubr v0.4.0 (Kassambara, 2020). We split the data by Phase 1 (comparison of ethanol treatments and DNA extraction protocols) and Phase 2 (comparison of types and amounts of starting material). Assignments were corrected: family and genera containing a single species were reassigned to that species, species were reassigned to domestic subspecies, and misassignments were corrected. Manual reassignment duplicated some assignments thus the read count data for these assignments were merged. Proportional read counts for each taxon were calculated from the total read counts per sample. Taxon-specific sequence thresholds (i.e. maximum sequence frequency for each taxon detected in process controls) were applied to the sponge samples to mitigate cross-contamination and false positives (Fig. S1). A second sequence threshold (0.01%) was applied to the sponge samples to mitigate low noise detections. Higher taxonomic assignments, iridescent catfish (PCR positive control), human, domestic species, and terrestrial species were then removed. Taxonomic assignments remaining in the refined dataset were predominantly of species resolution and considered true positives (Fig. S2).

#### 2.7.2 Data summaries

We examined overall fish and mammal taxon richness and read counts across samples processed with each ethanol treatment and DNA extraction protocol in Phase 1 (Fig. S3) and samples of each type and amount of starting material in Phase 2 (Fig. S4) for each sponge species. We also examined detection consistency across extraction replicates, in terms of taxon richness and number of replicates each taxon was detected in, for samples processed with each ethanol treatment and DNA extraction protocol in Phase 1 (Fig. S5) and samples of each type and amount of starting material in Phase 2 (Fig. S6) for each sponge species.

#### 2.7.2 Statistical analysis

We compared total DNA yield, post-PCR DNA concentration, taxon richness and proportional read counts of individual samples according to ethanol treatment and DNA extraction protocol for Phase 1, and according to type and amount of starting material for Phase 2. Only fish detections were used to compare extraction protocols and ethanol treatments in terms of taxon richness, proportional read counts, and community similarity as the metabarcoding primers used were designed to target fish. The data were not normally distributed and the assumptions of a two-way Analysis of Variance (ANOVA) were violated, thus statistical treatment followed Generalised Linear Models (GLMs) with Quasi-Poisson error family or Gaussian error family (for total DNA yield and post-PCR concentration), Poisson error family and the log-link function or negative binomial error family (for taxon richness), or binomial error family (for proportional read counts). If a significant effect was found using the *drop1* function from the package stats v3.6.3, multiple pairwise comparisons were performed using the *pairs* function from the package emmeans v1.7.2 (Lenth, 2022). Where assumptions of GLMs were violated, the non-parametric Kruskal-Wallis Test from the package stats was used, and multiple pairwise comparisons were performed using the non-parametric Dunn’s Test (p-values adjusted with the Benjamini-Hochberg method) from the package FSA v0.9.1 (Ogle et al., 2020). In Phase 1, data were split by ethanol treatment to compare extraction protocols and/or vice versa where significant effects were found using GLMs. In Phase 2, data were split by type of starting material to compare amounts of starting material and/or vice versa where significant effects were found using GLMs.

The package betapart v1.5.6 (Baselga & Orme, 2012) was used to estimate total beta diversity, partitioned by nestedness-resultant (i.e. community dissimilarity due to taxon subsets) and turnover (i.e. community dissimilarity due to taxon replacement), across samples in Phases 1 and 2 with the *beta.multi* function. Total beta diversity and its two components (Jaccard dissimilarity) were estimated for sponge samples in both phases using the *beta.pair* function. For total beta diversity and both partitions, we calculated the average distance of group members to the group centroid in multivariate space using the *betadisper* function in the package vegan v2.5-7 (Oksanen et al., 2019). This was done for each group of samples within ethanol treatments and DNA extraction protocols in Phase 1 and within types and amounts of starting material in Phase 2. The distances of group members to the group centroid were subjected to an ANOVA to test for homogeneity of multivariate dispersions (MVDISP), that is whether the dispersions of one or more groups were different, prior to permutational multivariate analysis of variance (PERMANOVA). Community dissimilarity for total beta diversity and both partitions was visualised with Non-metric Multidimensional Scaling (NMDS) using the *metaMDS* function with 1000 permutations, and tested statistically with PERMANOVA using the function *adonis* in the package vegan. The PERMANOVA included an interaction term between ethanol treatment and DNA extraction protocol for Phase 1 and an interaction term between type and amount of starting material for Phase 2. Pre-defined cut-off values were used for effect size, where PERMANOVA results were interpreted as moderate and strong effects if R^2^ > 0.09 and R^2^ > 0.25 respectively. These values are broadly equivalent to correlation coefficients of *r* = 0.3 and 0.5 which represent moderate and strong effects accordingly (Nakagawa & Cuthill, 2007; Macher et al., 2018).

## 3. Results

### 3.1 Sequencing performance

The Phase 1 library generated 3,757,770 raw sequence reads, of which 3,532,041 remained after quality control and merging. Following dereplication, denoising, and chimaera detection, 2,995,356 reads (average read count 31,201 per sample including controls) remained for taxonomic assignment. The Phase 2 library generated 3,005,701 raw sequence reads, of which 2,879,596 remained after quality control and merging. Following dereplication, denoising, and chimera detection, 2,170,282 reads (average read count 16,823 per sample including controls) remained for taxonomic assignment. For *L. chondrodes* samples, 85.05% of remaining reads were taxonomically assigned while 15.95% could not be assigned to any taxonomic rank. For *P. ventilabrum* and *V. pourtalesii* samples, 78.87% of remaining reads were taxonomically assigned, while 21.13% could not be assigned to any taxonomic rank. Contamination was restricted to human DNA in one extraction blank and one PCR negative control in Phase 1, but naturally occurring marine species were also detected in extraction blanks and PCR positive controls in Phase 2. Before dataset refinement, 29 taxa were identified in Phase 1 and 94 taxa were identified in Phase 2. After dataset refinement, 18 taxa were identified in Phase 1 and 49 taxa were identified in Phase 2 (Fig. S2).

For simplicity, we only report significant results from the final statistical analyses applied to the data. Full results from all statistical tests, including non-significant results and violations of model assumptions, are detailed in the Supporting Information. Statistical comparisons were not performed for DNA purity due to lack of variation in A260/A280 (Fig. 2c) and A260/A230 (Fig. 2d) ratios between ethanol treatments, extraction protocols, types of starting material, and amounts of starting material.

**Figure 2.**
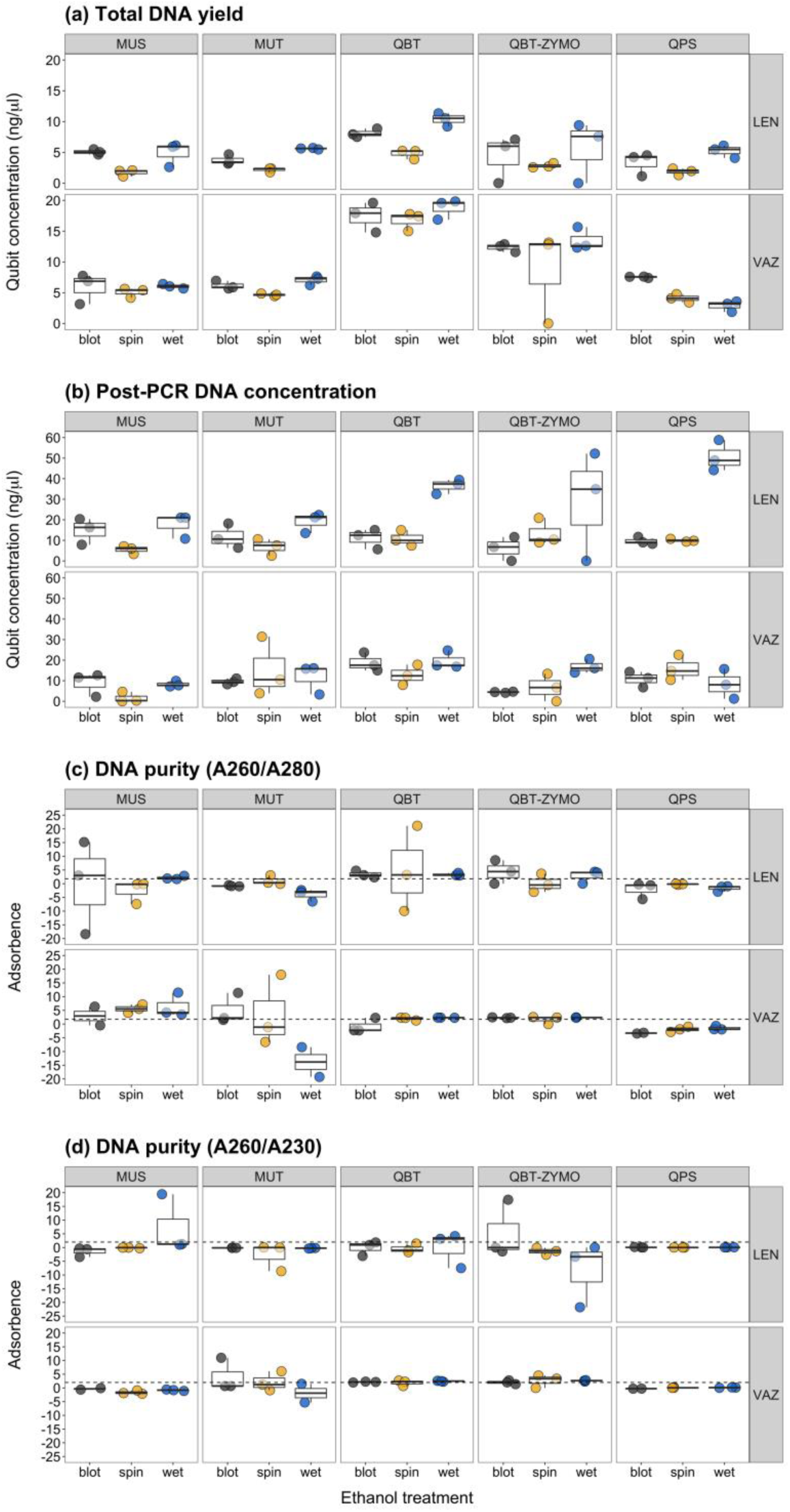
Box plots summarising **(a)** total DNA yield, **(b)** post-PCR DNA concentration, **(c)** A260/280 ratios, and **(d)** A260/A230 ratios for each sample from each sponge species processed with each ethanol treatment (blot, spin, wet) and each DNA extraction protocol (MUS, MUT, QBT, QBT-ZYMO and QPS). Abbreviations: *Lendenfeldia chondrodes* (LEN), *Vazella pourtalesii* (VAZ).

### 3.2 Phase 1: ethanol treatment and DNA extraction protocol

#### 3.2.1 Total DNA yield

For *L. chondrodes*, total DNA yield significantly differed between extraction protocols for the spin ethanol treatment (Kruskal-Wallis Test: H_4_ = 11.314, P = 0.023) and between ethanol treatments for the QBT (Kruskal-Wallis Test: H_2_ = 7.200, P = 0.027) and MUT (Kruskal-Wallis Test: H_2_ = 7.261, P = 0.027) protocols. For the spin treatment, none of the Dunn’s Test adjusted p-values were significant, but the largest differences in effect size were observed between the QBT and MUS protocols (Z = −2.749, P = 0.060), the QBT and MUT protocols (Z = −2.107, P = 0.117), and the QBT and QPS protocols (Z = 2.565, P = 0.052). For both the QBT and MUT protocols, the spin treatment produced lower total DNA yields than the wet treatment (QBT: Z = −2.683, P = 0.022; MUT: Z = −2.695, P = 0.021) (Fig. 2a). With *V. pourtalesii*, total DNA yield significantly differed between extraction protocols for the wet (Kruskal-Wallis Test: H_4_ = 13.233, P = 0.010) and blot (Kruskal-Wallis Test: H_4_ = 11.833, P = 0.019) ethanol treatments. The QBT protocol produced higher total DNA yields than the QPS protocol with the wet ethanol treatment (Dunn’s Test: Z = 3.286, P = 0.010) and the MUT protocol with the blot treatment (Dunn’s Test: Z = −2.921, P = 0.035) (Fig. 2a). For both sponges, the QBT protocol produced higher total DNA yields than other extraction protocols with the blot and wet ethanol treatments.

#### 3.2.2 Post-PCR DNA concentration

For *L. chondrodes*, ethanol treatment influenced post-PCR DNA concentration with the MUS protocol (Kruskal-Wallis Test: H_2_ = 6.006, P = 0.050), where the spin treatment produced lower post-PCR DNA concentrations than the wet treatment (Dunn’s Test: Z = −2.395, P = 0.050) (Fig. 2b). For *V. pourtalesii*, post-PCR DNA concentration was positively influenced by the MUT (GLM: 0.669 ± 0.327, t = 2.044, P = 0.048) and QBT (GLM: 1.005 ± 0.311, t = 3.229, P = 0.003) protocols, but the MUT protocol (12.13 ± 2.31 ng/μl) produced lower post-PCR DNA concentrations than the QBT protocol (16.97 2.74 ng/μl; pairwise comparison: Z = −3.229, P = 0.011) (Fig. 2b). Therefore, while the spin treatment tended to yield generally lower post-PCR DNA concentrations, differences did not appear to be substantial or consistent across sponges.

#### 3.2.3 Taxon richness

Taxon richness was visualised according to each extraction replicate of each sample (Fig. 3) and according to overall richness per sample (Fig. 4a), with the latter compared statistically. Only the MUS and QBT extraction protocols produced incidental mammal detections (Fig. 3). Using a GLM, neither extraction protocol or ethanol treatment were found to influence fish taxon richness for *L. chondrodes*, but model assumptions were met. For *V. pourtalesii*, no effect of extraction protocol on fish taxon richness was found using Kruskal-Wallis Test (Fig. 4a). For both sponges, neither extraction protocol nor ethanol treatment influenced fish taxon richness.

**Figure 3.**
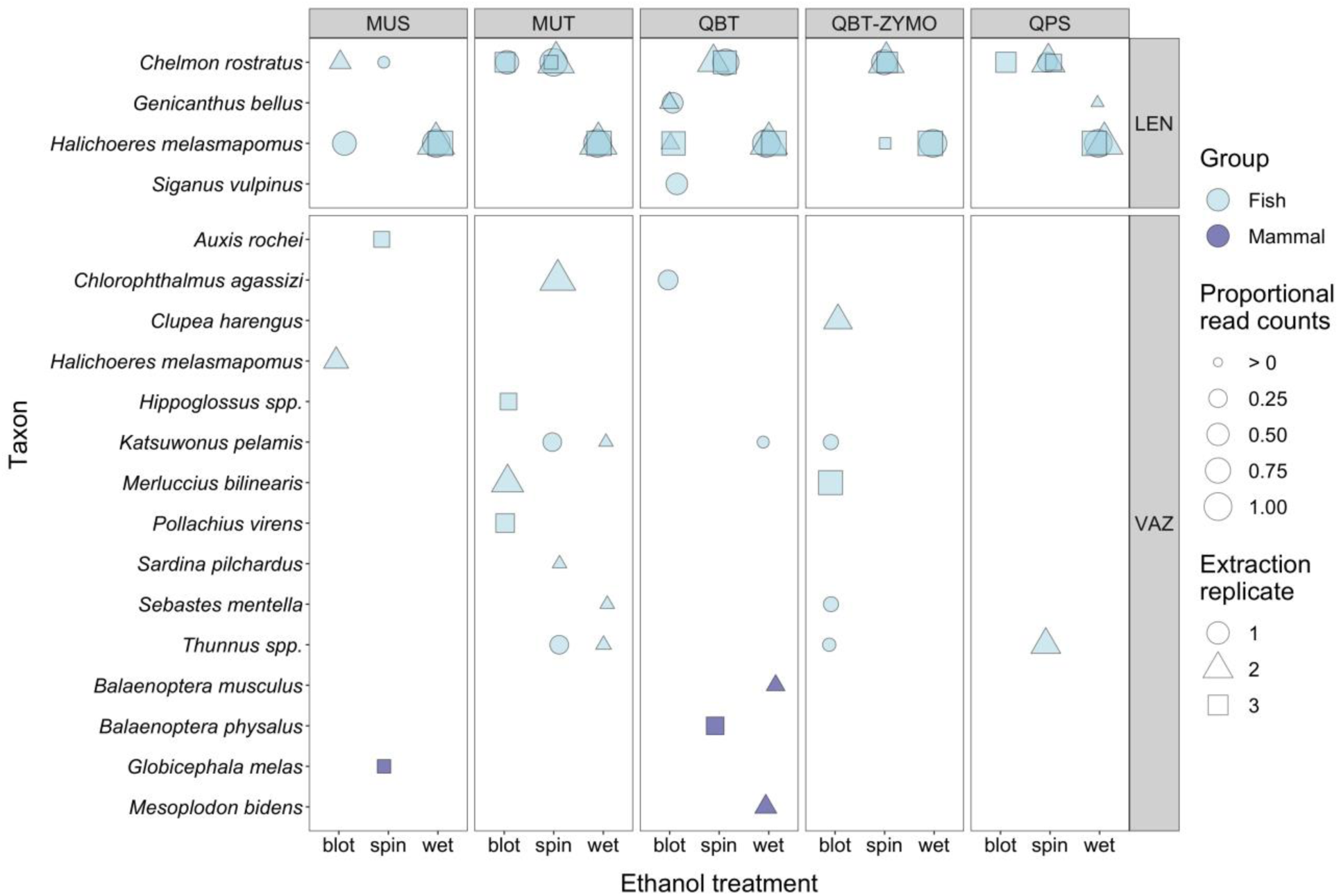
Bubble plot summarising proportional read counts for each taxon and detection consistency across extraction replicates for samples from each sponge species that were processed with each ethanol treatment (blot, spin, wet) and DNA extraction protocol (MUS, MUT, QBT, QBT-ZYMO and QPS). Abbreviations: *Lendenfeldia chondrodes* (LEN), *Vazella pourtalesii* (VAZ).

**Figure 4.**
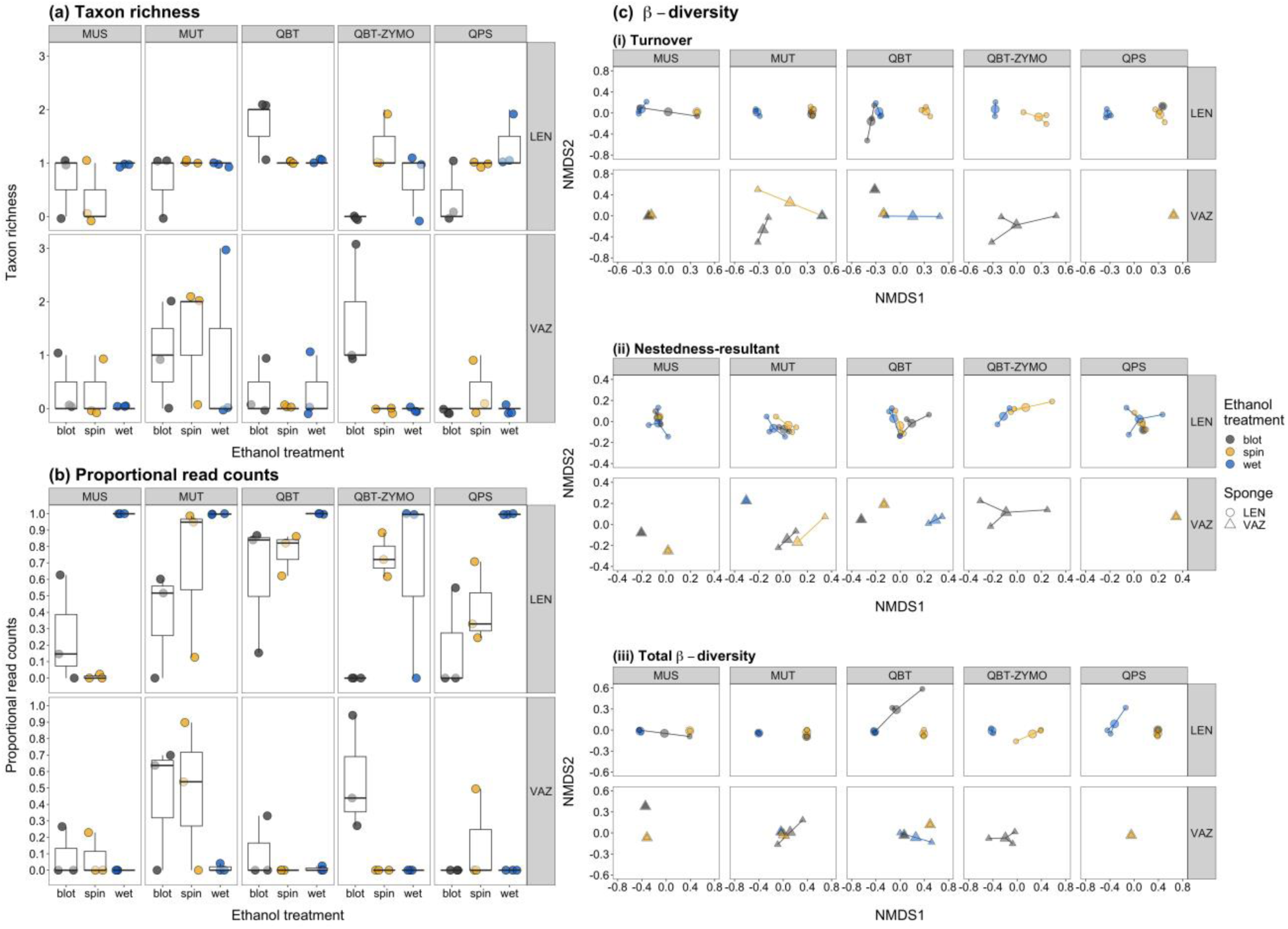
Differences in fish taxon richness **(a)**, proportional read counts **(b)** and community similarity **(c)**, including turnover **(i)** and nestedness-resultant **(ii)** components of total beta diversity **(iii)**, for samples processed with each ethanol treatment (blot, spin, wet) and each DNA extraction protocol (MUS, MUT, QBT, QBT-ZYMO and QPS). Abbreviations: *Lendenfeldia chondrodes* (LEN), *Vazella pourtalesii* (VAZ).

#### 3.2.4 Proportional read counts

Proportional read counts significantly differed between ethanol treatments with the MUS (Kruskal-Wallis Test: H_2_ = 6.161, P = 0.046) and QPS (Kruskal-Wallis Test: H_2_ = 6.006, P = 0.050) protocols for *L. chondrodes*. The wet treatment produced higher proportional read counts than the spin treatment with the MUS protocol (Dunn’s Test: Z = −2.426, P = 0.046), and than the blot treatment with the QPS protocol (Dunn’s Test: Z = −2.395, P = 0.050) (Fig. 4b). For *V. pourtalesii*, no effect of extraction protocol on proportional read counts was found, but proportional reads counts significantly differed between ethanol treatments with the QBT-ZYMO protocol (Kruskal-Wallis Test: H_2_ = 7.624, P = 0.022), where the blot treatment produced higher proportional read counts than the spin (Dunn’s Test: Z = 2.391, P = 0.025) or wet (Dunn’s Test: Z = 2.391, P = 0.050) treatments (Fig. 4b). Therefore, different extraction protocols and different ethanol treatments performed best in terms of proportional read counts for each sponge.

#### 3.2.5 Community composition

MVDISP differed between DNA extraction protocols for total beta diversity for *V. pourtalesii*, but not either partition for *V. pourtalesii* or *L. chondrodes* (Table 1). MVDISP differed between ethanol treatments for total beta diversity and the turnover component for *L. chondrodes*, and for the nestedness-resultant component of beta diversity for *V. pourtalesii*. DNA extraction protocol alone did not influence beta diversity for either sponge species (Table 2; Figs. 4ci-iii). Ethanol treatment had a strong positive influence on turnover and total beta diversity of communities detected from *L. chondrodes*, but did not influence beta diversity for *V. pourtalesii* (Table 2; Figs. 4ci-iii). DNA extraction protocol combined with ethanol treatment also had a moderate positive effect on turnover and total beta diversity for *L. chondrodes* (Table 2; Figs. 4ci-iii). Therefore, taxa detected from *L. chondrodes* with one ethanol treatment were replaced by different taxa detected with other ethanol treatments, although this finding should be interpreted with caution due to MVDISP heterogeneity.

**Table 1.**
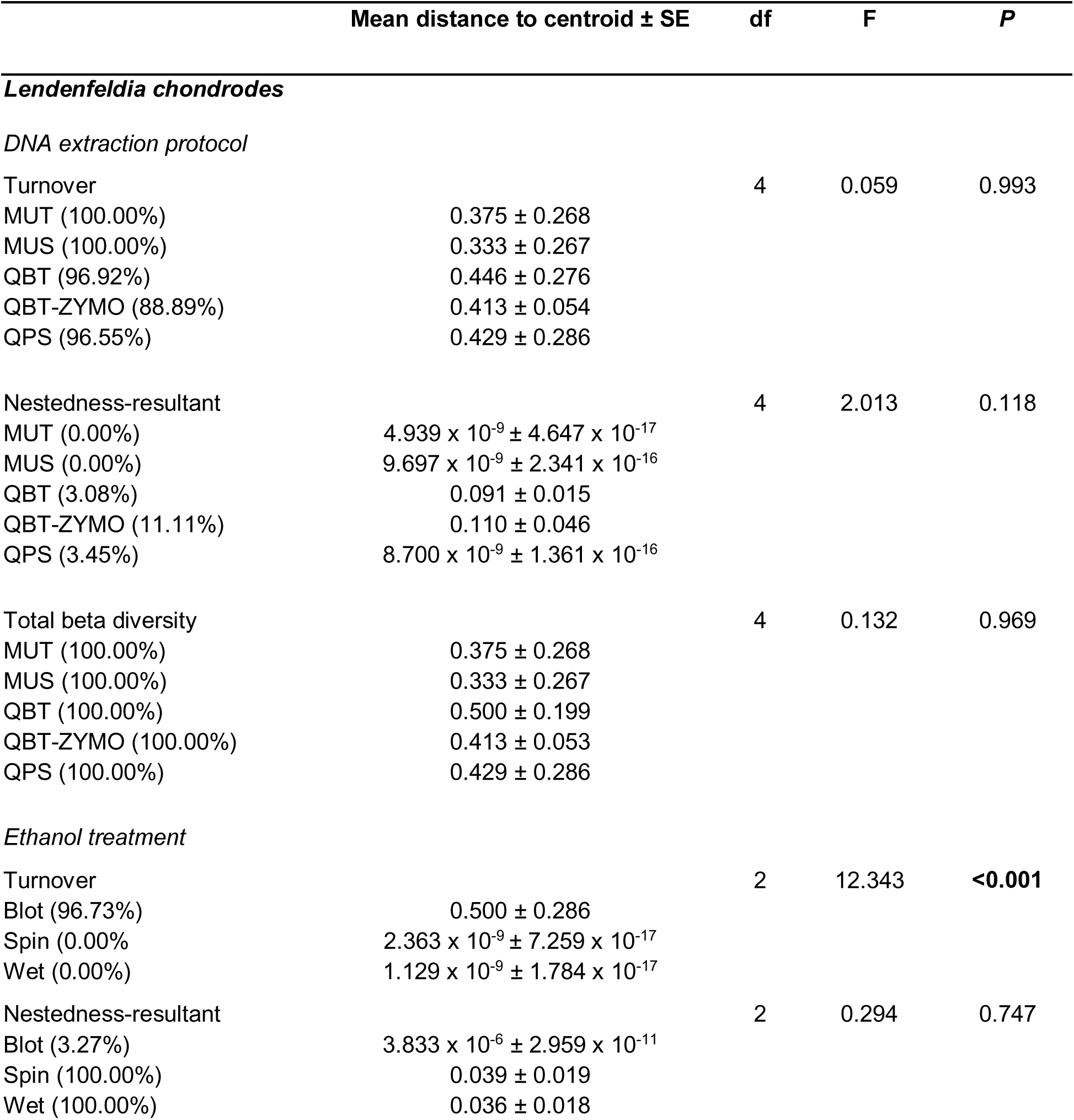
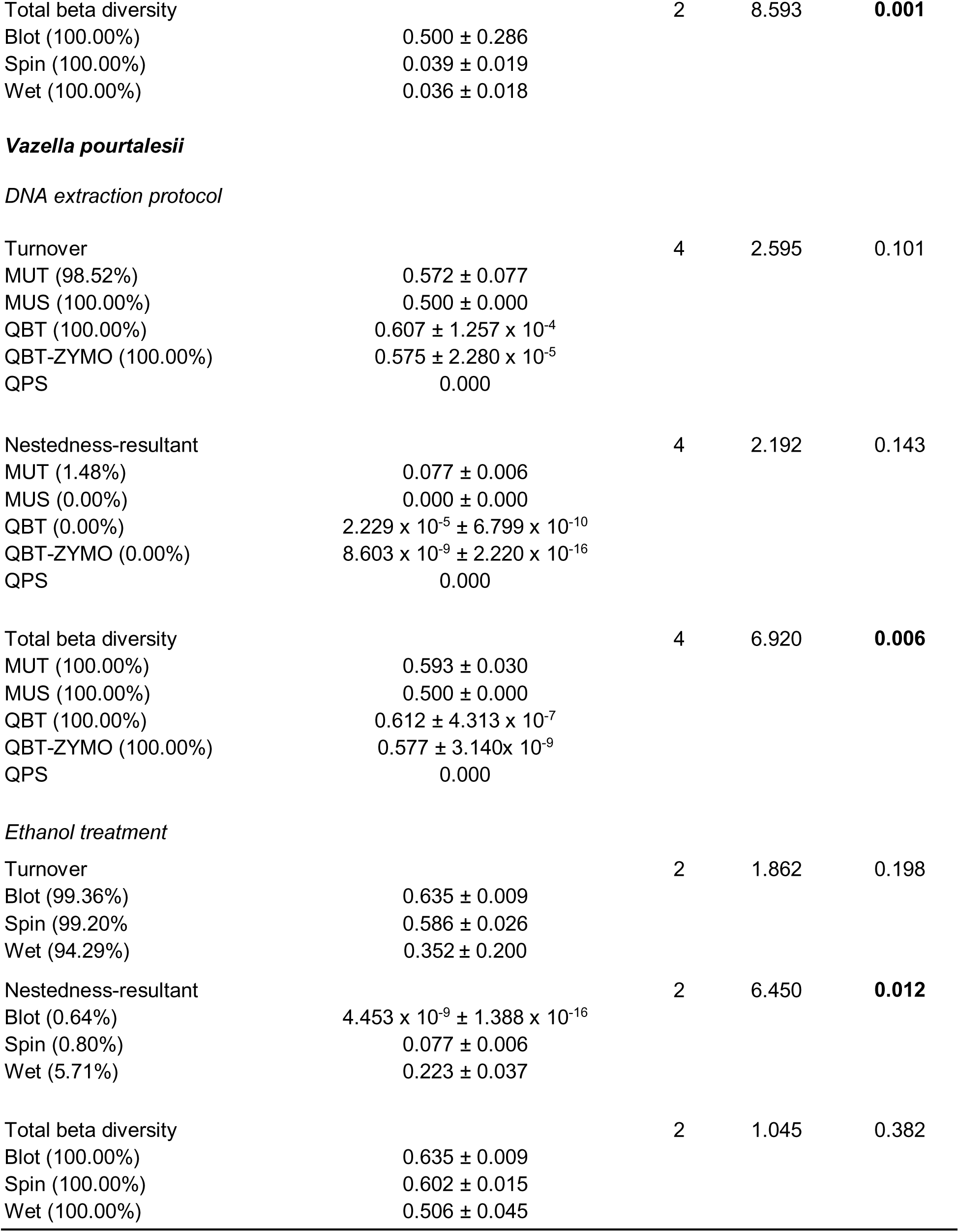
Summary of analyses statistically comparing homogeneity of multivariate dispersions (MVDISP) between the communities (ANOVA) produced by different extraction protocols and ethanol treatments. Relative contributions of taxon turnover and nestedness-resultant to total beta diversity (Jaccard dissimilarity) for each protocol and treatment are given in brackets.

**Table 2.** Results of PERMANOVA used to compare variation in community composition produced by different extraction protocols and ethanol treatments, individually and combined.

|  | df | F | R<sup>2</sup> | P | Stress |
| --- | --- | --- | --- | --- | --- |
| ***Lendenfeldia chondrodes*** |  |  |  |  | 0 |
| *Turnover* |  |  |  |  |  |
| DNA extraction protocol | 4 | 2.782 | 0.060 | 0.047 |  |
| Ethanol treatment | 2 | 67.309 | 0.720 | **0.001** |  |
| DNA extraction protocol :<br>Ethanol treatment | 7 | 2.883 | 0.108 | **0.022** |  |
| *Nestedness-resultant* |  |  |  |  | 0 |
| DNA extraction protocol | 4 | 0.670 | 0.122 | 0.570 |  |
| Ethanol treatment | 2 | -2.124 | -0.193 | 0.952 |  |
| DNA extraction protocol :<br>Ethanol treatment | 7 | 0.361 | 0.115 | 0.704 |  |
| *Total beta diversity* |  |  |  |  | 0 |
| DNA extraction protocol | 4 | 2.152 | 0.063 | 0.095 |  |
| Ethanol treatment | 2 | 46.617 | 0.677 | **0.001** |  |
| DNA extraction protocol :<br>Ethanol treatment | 7 | 2.132 | 0.108 | **0.049** |  |
| ***Vazella pourtalesii*** |  |  |  |  |  |
| *Turnover* |  |  |  |  | 0 |
| DNA extraction protocol | 4 | 0.733 | 0.234 | 0.948 |  |
| Ethanol treatment | 2 | 1.008 | 0.161 | 0.445 |  |
| DNA extraction protocol :<br>Ethanol treatment | 3 | 0.861 | 0.206 | 0.679 |  |
| *Nestedness-resultant* |  |  |  |  | 3.547 x 10<sup>-5</sup> |
| DNA extraction protocol | 4 | -2.911 x 10<sup>-15</sup> | 0.879 | 0.900 |  |
| Ethanol treatment | 2 | 2.083 x 10<sup>-15</sup> | -0.314 | 0.163 |  |
| DNA extraction protocol :<br>Ethanol treatment | 3 | -1.924 x 10<sup>-15</sup> | 0.436 | 0.819 |  |
| *Total beta diversity* |  |  |  |  | 0 |
| DNA extraction protocol | 4 | 0.814 | 0.252 | 0.872 |  |
| Ethanol treatment | 2 | 0.951 | 0.147 | 0.538 |  |
| DNA extraction protocol :<br>Ethanol treatment | 3 | 0.914 | 0.213 | 0.643 |  |

### 3.3 Phase 2: treatment and amount of sponge tissue

Based on Phase 1, blotting sponge tissue dry after ethanol preservation followed by DNA extraction with the MUT protocol appeared to be the best combination for extracting DNA from tissue of naturally occurring marine sponges like *V. pourtalesii*. Therefore, this procedure was selected for Phase 2.

#### 3.3.1 Total DNA yield

Using GLMs, amount of starting material influenced total DNA yield for both *P. ventilabrum* (F_2_ = 16.485, P < 0.001) and *V. pourtalesii* (F_2_ = 3.795, P = 0.030), but type of starting material influenced total DNA yield for *P. ventilabrum* only (F_1_ = 14.641, P < 0.001). Model assumptions were met for *P. ventilabrum*. Total DNA yield was higher using homogenised tissue (0.460 ± 0.120, t = 3.840, P < 0.001) and increased as the amount of starting material increased (amount 2: 0.451 ± 0.136, t = 3.321, P = 0.002; amount 3: 0.981 ± 0.165, t = 5.931, P < 0.001). Dried tissue (30.30 ± 2.81 ng/μl) produced lower total DNA yields than homogenate (48.00 ± 3.82 ng/μl; Z = −3.840, P < 0.001). Amount 3 (63.10 ± 7.96 ng/μl) produced higher total DNA yields than amount 2 (37.10 ± 3.18 ng/μl; Z = −3.474, P = 0.002) or amount 1 (23.70 ± 2.53 ng/μl; Z = −5.931, P < 0.001). Amount 1 also produced lower total DNA yields than amount 2 (Z = −3.321, P = 0.003) (Fig. 5a). Model assumptions were not met for *V. pourtalesii* thus Kruskal-Wallis Test was applied, but amount of starting material was not found to influence total DNA yield. For both sponges, amount of starting material appeared to influence total DNA yield, whereas type of starting material may have a more variable effect.

**Figure 5.**
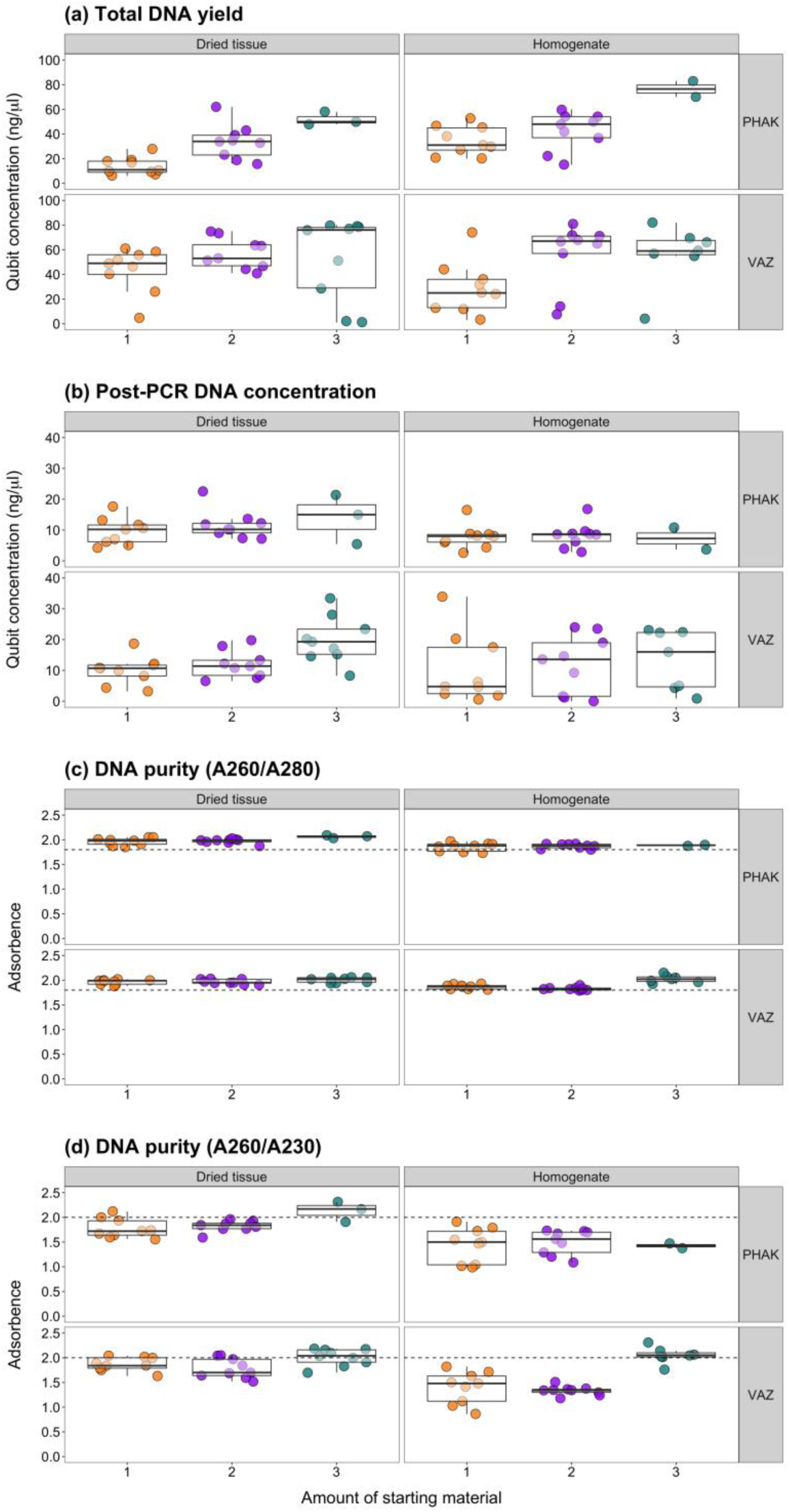
Box plots summarising **(a)** total DNA yield, **(b)** post-PCR DNA concentration, **(c)** A260/280 ratios, and **(d)** A260/A230 ratios for each sample from each sponge species processed using different types and amounts (1, 2, 3; see Figure 1 for volumes and weights used) of starting material. Abbreviations: *Phakellia ventilabrum* (PHAK), *Vazella pourtalesii* (VAZ).

#### 3.3.2 Post-PCR DNA concentration

Using GLMs, amount of starting material did not influence post-PCR DNA concentration for either *P. ventilabrum* or *V. pourtalesii*, and type of starting material influenced post-PCR DNA concentration for *P. ventilabrum* only (F_1_ = 5.054, P = 0.031). Model assumptions for *P. ventilabrum* were violated thus Kruskal-Wallis Test was applied, but no effect of type of starting material was found (Fig. 5b). For both sponges, neither type nor amount of starting material appeared to impact post-PCR DNA concentration.

#### 3.3.3 Taxon richness

Taxon richness was visualised according to each extraction replicate of each sample (Fig. 6) and according to overall richness per sample (Fig. 7a), with the latter compared statistically. *V. pourtalesii* produced more incidental mammal detections than *P. ventilabrum* (Fig. 6). Using GLMs, neither type or amount of starting material were found to influence fish taxon richness for *P. ventilabrum*, and type of starting material influenced fish taxon richness for *V. pourtalesii* only (χ_1_ = 6.451, P = 0.011). Type of starting material had a positive influence on taxon richness (0.476 ± 0.190, t = 2.514, P = 0.012), where taxon richness was higher using homogenate (2.780 ± 0.334) than dried tissue (1.720 ± 0.253; Z = −2.514, P = 0.012) (Fig. 6a). Therefore, amount of starting material did not influence fish taxon richness for either sponge, but type of starting material may be an important consideration for some sponges.

**Figure 6.**
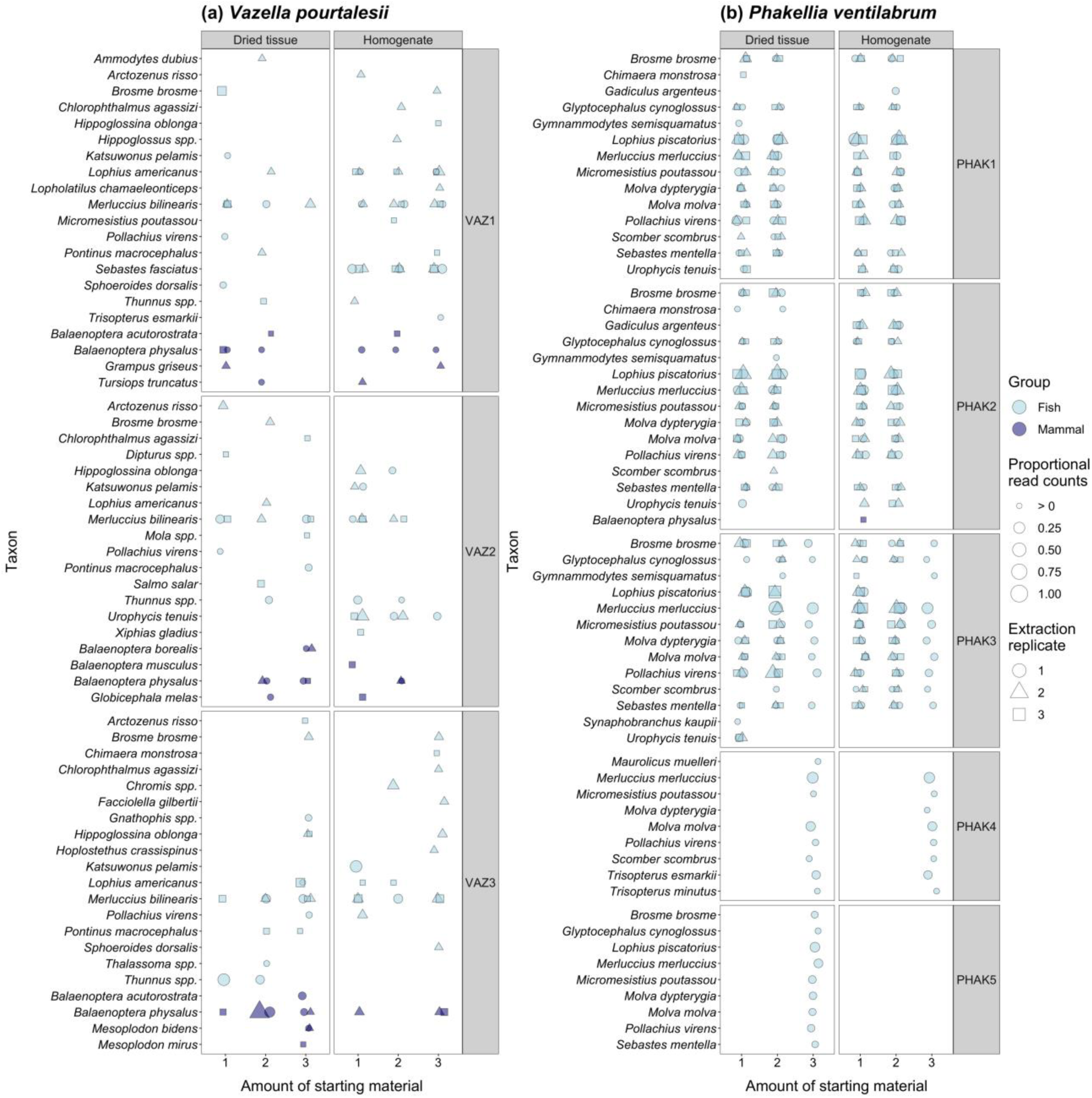
Bubble plots summarising proportional read counts for each taxon and detection consistency across extraction replicates for different types and amounts (1, 2, 3; see Figure 1 for volumes and weights used) of starting material from **(a)** *Vazella pourtalesii* (VAZ) and **(b)** *Phakellia ventilabrum* (PHAK).

**Figure 7.**
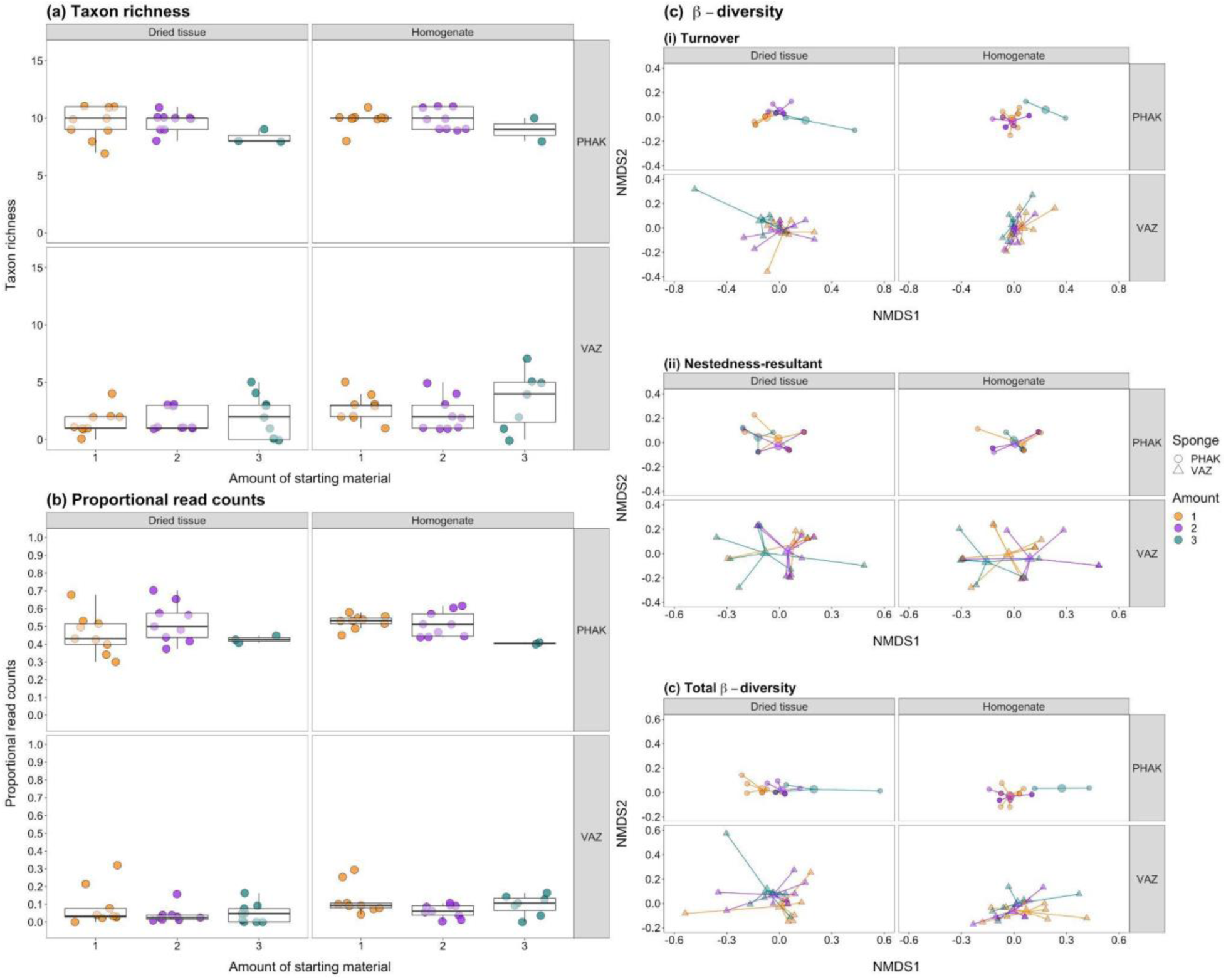
Differences in taxon richness **(a)**, proportional read counts **(b)** and community similarity **(c)**, including turnover **(i)** and nestedness-resultant **(ii)** components of total beta diversity **(iii)**, for fish detected in samples of each type and amount (1, 2, 3; see Figure 1 for volumes and weights used) of starting material from each sponge species. Abbreviations: *Phakellia ventilabrum* (PHAK), *Vazella pourtalesii* (VAZ).

#### 3.3.4 Proportional read counts

Using GLMs, neither type or amount of starting material were found to influence fish proportional read counts for *P. ventilabrum*, and only amount of starting material influenced fish proportional read counts for *V. pourtalesii* (F_2_ = 3.266, P = 0.047). Model assumptions for *V. pourtalesii* were violated thus Kruskal-Wallis Test was applied, but no effect of amount of starting material was found (Fig. 6b). For both sponges, there was no clear effect of type or amount of starting material on proportional read counts.

#### 3.3.5 Community composition

MVDISP did not differ between types of starting material for total beta diversity or its partitions for either sponge. MVDISP did not differ between amounts of starting material for total beta diversity or its partitions for *V. pourtalesii*, but did differ between amounts for total beta diversity and turnover for *P. ventilabrum* (Table 3). For *P. ventilabrum*, amount of starting material influenced turnover and total beta diversity, but type of starting material did not influence beta diversity (Table 4; Figs. 7ci-iii). For *V. pourtalesii*, type of starting material influenced turnover and total beta diversity, but amount of starting material did not influence beta diversity (Table 4; Figs. 7ci-iii). Therefore, taxa detected with one type or amount of starting material were replaced by different taxa detected with the other type (*V. pourtalesii*) or amounts (*P. ventilabrum*) of starting material, although the latter should be interpreted with caution due to MVDISP heterogeneity.

**Table 3.** Results of ANOVA used to compare homogeneity of multivariate dispersions (MVDISP) between the communities produced by different types and amounts of starting material. Relative contributions of taxon turnover and nestedness-resultant to total beta diversity (Jaccard dissimilarity) for each type and amount of starting material are given in brackets.

| | Mean distance to centroid ± SE | df | F | P |
| --- | --- | --- | --- | --- |
| ***Phakellia ventilabrum*** |  |  |  |  |
| *Type of starting material* |  |  |  |  |
| Turnover |  | 1 | 0.172 | 0.681 |
| Dried tissue (85.02%) | 0.115 ± 0.021 | | | |
| Homogenate (88.16%) | 0.131 ± 0.010 | | | |
| Nestedness-resultant |  | 1 | 1.729 | 0.196 |
| Dried tissue (14.98%) | 0.074 ± 0.004 | | | |
| Homogenate (11.84%) | 0.051 ± 0.002 | | | |
| Total beta diversity |  | 1 | 0.059 | 0.809 |
| Dried tissue (100.00%) | 0.165 ± 0.026 | | | |
| Homogenate (100.00%) | 0.175 ± 0.011 | | | |
| *Amount of starting material* |  |  |  |  |
| Turnover |  | 2 | 6.141 | **0.005** |
| 1 (79.35%) | 0.106 ± 0.005 | | | |
| 2 (79.46%) | 0.096 ± 0.004 | | | |
| 3 (92.15%) | 0.247 ± 0.038 | | | |
| Nestedness-resultant |  | 2 | 0.425 | 0.657 |
| 1 (20.65%) | 0.073 ± 0.005 | | | |
| 2 (20.54%) | 0.059 ± 0.002 | | | |
| 3 (7.85%) | 0.049 ± 0.002 | | | |
| Total beta diversity |  | 2 | 4.898 | **0.013** |
| 1 (100.00%) | 0.153 ± 0.009 | | | |
| 2 (100.00%) | 0.137 ± 0.005 | | | |
| 3 (100.00%) | 0.287 ± 0.030 | | | |
| ***Vazella pourtalesii*** |  |  |  |  |
| *Type of starting material* |  |  |  |  |
| Turnover |  | 1 | 0.327 | 0.570 |
| Dried tissue (97.03%) | 0.489 ± 0.059 | | | |
| Homogenate (96.63%) | 0.451 ± 0.046 | | | |
| Nestedness-resultant |  | 1 | 0.696 | 0.409 |
| Dried tissue (2.97%) | 0.146 ± 0.019 |  |  |  |
| Homogenate (3.37%) | 0.186 ± 0.035 |  |  |  |
| Total beta diversity |  | 1 | 0.143 | 0.707 |
| Dried tissue (100.00%) | 0.569 ± 0.024 |  |  |  |
| Homogenate (100.00%) | 0.554 ± 0.014 |  |  |  |
| *Amount of starting material* |  |  |  |  |
| Turnover |  | 2 | 0.165 | 0.849 |
| 1 (96.29%) | 0.513 ± 0.025 |  |  |  |
| 2 (97.05%) | 0.477 ± 0.055 |  |  |  |
| 3 (95.02%) | 0.480 ± 0.044 |  |  |  |
| Nestedness-resultant |  | 2 | 0.645 | 0.530 |
| 1 (3.71%) | 0.132 ± 0.010 |  |  |  |
| 2 (2.95%) | 0.177 ± 0.030 |  |  |  |
| 3 (4.98%) | 0.115 ± 0.037 |  |  |  |
| Total beta diversity |  | 2 | 0.022 | 0.978 |
| 1 (100.00%) | 0.569 ± 0.015 |  |  |  |
| 2 (100.00%) | 0.563 ± 0.021 |  |  |  |
| 3 (100.00%) | 0.559 ± 0.012 |  |  |  |

**Table 4.** Results of PERMANOVA used to compare variation in community composition according to type and amount of starting material, individually and combined.

|  | df | F | R<sup>2</sup> | P | Stress |
| --- | --- | --- | --- | --- | --- |
| ***Phakellia ventilabrum*** |  |  |  |  |  |
| *Turnover* |  |  |  |  | 0.065 |
| Type of starting material | 1 | 1.917 | 0.034 | 0.165 |  |
| Amount of starting material | 2 | 7.513 | 0.269 | **0.002** |  |
| Type of starting material : | 2 | 1.981 | 0.071 | 0.140 |  |
| Amount of starting material |  |  |  |  |  |
| *Nestedness-resultant* |  |  |  |  | 0.045 |
| Type of starting material | 1 | 1.941 | 0.054 | 0.178 |  |
| Amount of starting material | 2 | 0.385 | 0.021 | 0.689 |  |
| Type of starting material : | 2 | -0.756 | -0.042 | 0.991 |  |
| Amount of starting material |  |  |  |  |  |
| *Total beta diversity* |  |  |  |  | 0.075 |
| Type of starting material | 1 | 1.323 | 0.026 | 0.299 |  |
| Amount of starting material | 2 | 5.571 | 0.220 | **0.001** |  |
| Type of starting material : | 2 | 1.638 | 0.065 | 0.152 |  |
| Amount of starting material |  |  |  |  |  |
| ***Vazella pourtalesii*** |  |  |  |  |  |
| *Turnover* |  |  |  |  | 0.119 |
| Type of starting material | 1 | 3.058 | 0.067 | **0.011** |  |
| Amount of starting material | 2 | 0.664 | 0.029 | 0.745 |  |
| Type of starting material : | 2 | 0.507 | 0.022 | 0.863 |  |
| Amount of starting material |  |  |  |  |  |
| *Nestedness-resultant* |  |  |  |  | 0.189 |
| Type of starting material | 1 | -6.207 | -0.145 | 0.998 |  |
| Amount of starting material | 2 | 2.410 | 0.113 | 0.232 |  |
| Type of starting material : | 2 | 2.105 | 0.098 | 0.259 |  |
| Amount of starting material |  |  |  |  |  |
| *Total beta diversity* |  |  |  |  | 0.113 |
| Type of starting material | 1 | 2.055 | 0.045 | **0.017** |  |
| Amount of starting material | 2 | 1.026 | 0.045 | 0.410 |  |
| Type of starting material : | 2 | 0.751 | 0.033 | 0.841 |  |
| Amount of starting material |  |  |  |  |  |

## 4. Discussion

We have illustrated how the performance of sponge nsDNA metabarcoding can be affected by DNA isolation procedures, specifically treatment to remove residual ethanol prior to DNA extraction, DNA extraction protocol, type of starting material, and amount of starting material. These effects can also be sponge-specific. Our experiments have provided an optimised DNA isolation procedure that will inform the development and application of sponge natural sampler DNA metabarcoding for aquatic biodiversity assessment.

### 4.1 Phase 1: ethanol treatment and DNA extraction protocol

The QBT and MUT extraction protocols outperformed others for different ethanol treatments of sponge tissue in terms of total DNA yield and post-PCR DNA concentration. Lower DNA purity was observed with the MUT and QBT-ZYMO protocols for the wet ethanol treatments of *V. pourtalesii* tissue and *L. chondrodes* tissue respectively, but otherwise extraction protocols produced consistent purity ratios. Extraction protocol was not found to influence fish taxon richness, proportional read counts or community composition (arguably the most important extraction success criteria) for either sponge species.

The spin ethanol treatment produced lower total DNA yields and post-PCR DNA concentrations with certain extraction protocols for *L. chondrodes*, but no differences were observed between ethanol treatments for *V. pourtalesii*. Ethanol treatment did not influence fish taxon richness for either sponge species, but did influence fish proportional read counts with certain extraction protocols, where the wet or blot treatments produced higher fish proportional read counts than the spin treatment. Ethanol treatment influenced community composition of *L. chondrodes* tissue but not *V. pourtalesii* tissue, where different taxa were detected in *L. chondrodes* tissue with different ethanol treatments.

DNA extraction protocols have been found to influence the performance of metabarcoding across sample types and habitats (Deiner et al., 2018; Jeunen et al., 2019; Johnson et al., 2019; Geraldi et al., 2019; Sanches & Schreier, 2020), thus it was not surprising that this also turned out to be the case for sponge tissue. All extraction protocols produced some biodiversity information, but some protocols were more effective than others, which has implications for detection of rare and/or low density species in diverse and complex marine systems using sponge natural sampler DNA metabarcoding. Our findings echo those of Sellers et al. (2018), where the Mu-DNA protocols were comparable to or outperformed commercial kits, and other studies which found the QBT protocol produced higher DNA yield than other commercial kits and unbranded extraction protocols (Hinlo et al., 2017; Deiner et al., 2018; Jeunen et al., 2019). Mu-DNA is advantageous due to adaptability and cost-efficiency. Although an inhibitor removal step is not included in the tissue protocol published in Sellers et al. (2018), it can be added from the soil or water protocols, which is desirable for obtaining high DNA yields whilst removing inhibitors (Jeunen et al., 2019). A QBT extraction costs four times as much as a MUT extraction (Sellers et al., 2018), and requires a commercial post-extraction inhibitor removal kit (e.g. Zymo Research) to remove potential PCR inhibitors. Reduced cost means that the Mu-DNA protocols are more scalable for large projects and buffer volumes can be adjusted proportionately to the amount of starting material without quickly depleting budgets.

Residual ethanol in tissue can retain unique DNA that may leech from tissue into ethanol (Zizka et al., 2018; Marquina et al., 2019; Jeunen et al., 2019; Derycke et al., 2021; Persaud et al., 2021; Wang et al., 2021), but also react with buffers used for DNA extraction and/or inhibit PCR amplification downstream (Schrader et al., 2012). Our DNA concentration results seem to indicate that residual ethanol contained DNA unique and/or additional to that present in sponge tissue (at least for *L. chondrodes*) as higher total DNA yields and post-PCR DNA concentrations were observed with the wet ethanol treatment. However, this did not translate to higher fish taxon richness, and conflicting effects were found for fish proportional read counts from each sponge species, where the wet ethanol treatment generated higher proportional read counts for *L. chondrodes* and the blot ethanol treatment produced higher proportional read counts for *V. pourtalesii*. Beta diversity analyses also indicated that different taxa were detected using different ethanol treatments for *L. chondrodes*. This reaffirms findings of studies on bulk invertebrate samples (Zizka et al., 2018; Derycke et al., 2021; Persaud et al., 2021; Wang et al., 2021) and marine sponges (Jeunen et al., 2019) where eDNA from ethanol preservative detected different taxa to DNA from tissue, although eDNA from ethanol preservative generally detected more taxa than bulk invertebrate tissue but less taxa than marine sponge tissue.

Sponge species, extraction protocol and ethanol treatments appear to have specific interactions and effects on total DNA yield, DNA purity, target DNA concentration, taxon recovery, and sequence generation. This has been observed for aquatic eDNA in terms of filter material and extraction protocol (Deiner et al., 2017), although other studies have suggested that these aspects of the eDNA workflow do not interact with each other (Sanches & Schreier, 2020). A more elaborate experiment incorporating numerous individuals of different sponge species would be required to test this hypothesis. Based on our results, we recommend that these future studies focus on comparing tissue-based extraction protocols and DNA extraction from ethanol alone versus sponge tissue alone versus ethanol and sponge tissue combined. For the purposes of experimental Phase 2 in the present study, blotting sponge tissue dry after ethanol preservation followed by DNA extraction with the MUT protocol appeared to be the best combination for extracting DNA from tissue of naturally occurring marine sponges like *V. pourtalesii*.

### 4.2 Phase 2: treatment and amount of sponge tissue

Homogenised tissue produced higher total DNA yields for *P. ventilabrum* tissue but not *V. pourtalesii* tissue. DNA from homogenised tissue from both sponge species was of lower purity. Type of starting material did not influence post-PCR DNA concentration for either sponge species. Homogenised tissue resulted in higher fish taxon richness for *V. pourtalesii*, although not *P. ventilabrum*, and type of starting material did not influence fish proportional read counts for either sponge species. Different communities were observed using different types of starting material for *V. pourtalesii*, but community composition from homogenate and dried tissue was highly consistent for *P. ventilabrum*. Larger amounts of starting material produced higher total DNA yields for *P. ventilabrum* tissue, but not *V. pourtalesii* tissue. DNA purity was consistent across different amounts of starting material for both sponge species. Amount of starting material did not influence post-PCR DNA concentration, fish taxon richness or fish proportional read counts for either sponge species. However, different communities were observed using different amounts of starting material for *P. ventilabrum*, but not for *V. pourtalesii*.

Homogenisation is often used with complex sample types to account for uneven distribution of organisms/DNA throughout the sample by facilitating the release of DNA via bead beating, blending or vortexing, but introduces contamination risk via more handling of samples and reusable equipment (Gosselin et al., 2017; Marquina et al., 2019; Pereira-da-Conceicoa et al., 2020; Hestetun et al., 2021; Zizka et al., 2022). Until now, the distribution of eDNA in marine sponge tissue and whether homogenisation was required was unknown. Our results suggest that eDNA has a relatively even distribution in sponge tissue. Type of starting material did not influence fish taxon richness for *P. ventilabrum* or fish proportional read counts for either sponge species. Although higher taxon richness was observed with homogenised *V. pourtalesii* tissue, the gains were marginal compared to dried *V. pourtalesii* tissue. A homogenous distribution of eDNA in sponge tissue offers additional advantages to using these natural samplers for marine biodiversity assessment over aquatic eDNA which can exhibit substantial spatial heterogeneity (Bruce et al., 2021). Dried tissue appears to be sufficient for taxon recovery and would avoid the contamination risk associated with homogenisation. However, the accumulation and degradation rates of eDNA in sponge tissues still require investigation to quantify temporal variation in eDNA signals from these natural samplers compared to aquatic eDNA (Jeunen et al., 2021), which may differ by target taxa and habitat (Harrison et al., 2019). This has recently been done in an artificial setting (Cai et al., 2022), but must also be investigated in natural habitats.

Total DNA yield increased as volume of homogenate and weight of dry tissue increased. This relationship was expected (bar the risk of introducing inhibitors by using too much tissue for DNA extraction) as using more starting material was found to improve detection success for metabarcoding of bulk invertebrate (Zizka et al., 2022) and soil (Kirse et al., 2021) samples. However, more starting material did not result in higher fish taxon recovery or fish proportional read counts, which were consistent across volumes of homogenate and weights of dried tissue. Different amounts of starting material also produced similar communities for *V. pourtalesii*, albeit different communities were observed with different amounts of *P. ventilabrum* tissue. Therefore, there seems to be little benefit to using more starting material in sponge natural sampler DNA metabarcoding as more complex extraction procedures are required to handle larger quantities of tissue. Instead, it may be more advantageous to invest effort in performing more extraction replicates with smaller amounts of starting material (Lanzén et al., 2017; Hestetun et al., 2021).

### 4.3 Future directions

Our findings have advanced the field of sponge nsDNA metabarcoding and emphasise the importance of sequencing samples used in comparisons of DNA extraction procedures rather than relying only on DNA concentration or purity values and gel images. Nevertheless, a number of unanswered questions remain. The sample size used in our study for each sponge, ethanol treatment, and DNA extraction protocol as well as type and amount of starting material was small due to the various permutations on DNA extraction methods being investigated and the available budget. Thousands of sponges occupy marine systems at every depth and latitude across the globe (Morganti et al., 2021). They can vary in shape, size, and hardness giving rise to variable characteristics, including height, diameter, depth, growth form (free-standing, encrusting, tubular, massive), substrate (hard rock, soft sediment, animal, floating debris), spicules (spongin fibre, calcium carbonate, silica), colour, symmetry (radial symmetry or asymmetrical), body plan (Asconoid, Syconoid, Leuconoid), and number and cross-sectional area of oscula. They also differ in terms of reproduction (sexual or asexual), feeding (carnivorous or herbivorous), defence (toxicity or boring activity), disease (fungi, viruses, cyanobacteria), and symbionts (fish, shrimps, plants, bacteria, algae) (Kahn et al., 2015; Kandler et al., 2019; Morganti et al., 2021). Only by pooling empirical natural sampler DNA-inferred taxon inventories from a variety of wild sponges in natural settings will it be possible to gauge a more comprehensive understanding of the robustness and reliability of this approach across habitats and geographical regions.

Different sponges may not have the same pumping and filtration efficiency or even have viable tissue for DNA extraction due to microbial loads and PCR inhibitors (Cai et al., 2022). Therefore, future research must identify the best natural samplers in both benthic and pelagic habitats from those sponges that are accessible and can withstand repeat sampling as well as understand how different sponge characteristics influence eDNA recovery (Mariani et al., 2019; Jeunen et al., 2021). The spatial and temporal resolution of sponge natural sampler DNA under different environmental conditions and across seasons must be investigated as we do not know the spatial extent or recency of eDNA trapped in sponge tissue (Turon et al., 2020) and whether eDNA in sponge tissue is affected by the same factors that hasten degradation of aquatic eDNA (e.g. temperature, pH, salinity, microbial activity), but these factors are known to influence the pumping and filtration action of sponges (Gökalp et al., 2020; Morganti et al., 2021). It is also undetermined whether sponges exhibit selectivity for eDNA particles of different sizes akin to filter membranes used for aquatic eDNA (Turner et al., 2014) and the sampling effort (whether replicates from the same individual or replicates from different individuals) required for sponges to achieve set species detection probabilities (Turon et al., 2020).

In addition to these biological considerations, there are further technical considerations for sponge natural sampler DNA metabarcoding. Many species were only detected in one extraction replicate performed on tissue from wild *V. pourtalesii* specimens, emphasising the potential stochasticity of sponge tissue and need for extraction replicates, similar to sediment (Lanzén et al., 2017; Hestetun et al., 2021) and bulk invertebrate samples (Elbrecht & Steinke, 2018). Marker and primer choice is crucial to metabarcoding success. Existing sponge natural sampler DNA studies have focused on 12S and 16S rRNA primers for the detection of fish (Mariani et al., 2019; Turon et al., 2020; Jeunen et al., 2021), but as future studies diverge and other markers are explored, primers that minimise non-target amplification (Collins et al., 2019) and blocking primers (Boessenkool et al., 2012) to prevent amplification of DNA from the sponge itself could become essential (Jeunen et al., 2021).

Finally, sequencing depth may play an important role in detection of low concentration eDNA signals in sponge tissue, whether these represent species that are rare, small or release less DNA into the environment. Very few studies have compared sequencing platforms for metabarcoding, but comparisons have been made for the popular Illumina MiSeq with the larger Illumina NovaSeq (Singer et al., 2019) and with the smaller Illumina iSeq 100 (Nakao et al., 2021). The Illumina NovaSeq detected more species from seawater than the Illumina MiSeq, and higher sequencing depth was also found to improve species detection from sediment (Lanzén et al., 2017) and faecal (Alberdi et al., 2018) samples with metabarcoding. However, species detection rates and community composition were highly similar between the Illumina MiSeq and Illumina iSeq 100 (Nakao et al., 2021). Given these disparities, future studies should carefully consider sequencing depth for natural sampler DNA metabarcoding when testing the performance of new natural samplers and addressing the outstanding questions identified here.

These biological and technical considerations are well-exemplified by *L. chondrodes* in our study which detected 4 of 25 (after dataset refinement) or 11 of 25 (before dataset refinement) species present in the aquarium tank. The sponge individuals were relatively young, having regrown over a few weeks after attempts to remove the sponge from the aquarium tank. This should have been enough time for eDNA to accumulate in the tissues of these sponges (Cai et al., 2022), but they may not have reached a large enough size to adequately pump and filter eDNA from the volume of water held in the tank. Furthermore, it is unknown whether bacterial symbionts, such as those observed in *L. chondrodes* (Galitz et al., 2018), influence the pumping and filtration efficiency of sponges, although results from Cai et al. (2022) appear to suggest that symbiotic load may influence eDNA capture and persistence in sponge tissue. The BLAST identity for taxonomic assignment also had to be reduced to 80% in order to recover more species known to be present in the aquarium tank. This suggests that the primers used may have insufficient taxonomic resolution to discriminate between tropical fish species, namely members of the families Acanthuridae, Chaetodontidae, Labridae, Pomacanthidae, Serranidae and Siganidae.

## 5. Conclusions

Our study represents another step forward in the development of sponge natural sampler DNA metabarcoding as a means to assess aquatic biodiversity, and highlights how methodological choices (specifically DNA isolation procedures) can influence the taxa and subsequent communities recovered using this approach. Based on our results, we recommend blotting sponge tissue dry to remove ethanol preservative prior to extraction with a tissue-based protocol, whether commercial or non-commercial. However, we note that more extensive comparisons of eDNA in ethanol preservative only, sponge tissue only, and ethanol and sponge tissue combined extracted with tissue-based protocols would be worthwhile. Homogenisation does not seem to be necessary for sponges, eliminating a time-consuming step that presents a contamination risk. Amount of starting material did not substantially influence taxon recovery or community composition, but further comparisons of multiple extraction replicates with small amounts of sponge tissue versus a single extraction with a large amount of tissue are needed to determine whether time is better invested in simpler but more numerous extractions or more complex but fewer extractions. With these additional comparative studies, we will have a comprehensive understanding of the optimal DNA isolation procedure for sponge nsDNA metabarcoding which future investigations, whether of a biological or technical nature, can draw upon to advance this growing field.

## Acknowledgements

This study was funded by the UK Natural Environment Research Council grant NE/T007028/1. A.R. was also supported by the Spanish Ministry of Science and Innovation grant (RYC2018-024247-I) and CSIC intramural grant (202030E006). M.B.A. was supported by the National Agency for Research and Development (ANID), fellowship program Postdoctorado en el extranjero/2019 - 74200143.

## Author Contributions

S.M. and L.R.H. designed the study; J.C., A.R., and B.M. facilitated sample collection; L.R.H, E.F.N. and A.C. performed the laboratory work with guidance from G.S., M.B.A. and A.R.; G.S. performed the bioinformatic analyses; S.M. and A.R. obtained the funding; L.R.H. analysed the data and wrote the manuscript, which all authors revised.

## Data Accessibility

Raw sequence reads have been archived on the NCBI Sequence Read Archive (BioProject: PRJNA854174; BioSamples: SAMN29421799 - SAMN29421894 [Phase 1] and SAMN29444657 - SAMN29444784 [Phase 2]; SRA accessions: SRR19906974 - SRR19907069 [Phase 1] and SRR19912485 - SRR19912613 [Phase 2]). Scripts and corresponding data have been deposited in a dedicated Zenodo repository (https://doi.org/10.5281/zenodo.7264066).

