## Supporting Information for "Optimised DNA isolation from marine sponges for natural sampler DNA (nsDNA) metabarcoding"

### Materials and Methods

#### Workspace and decontamination procedures

A unidirectional workflow was used for sample processing. Sponge tissue was stored and handled in a laboratory dedicated to the processing of environmental samples with low DNA concentrations. Users of this laboratory put on a disposable hooded bodysuit, disposable shoe covers, two pairs of disposable gloves secured at the wrist with masking tape, and a face mask in a separate entry room before entering the main laboratory.

All bench spaces and equipment were sterilised before and after use by wiping with fresh paper towel and 10% v/v bleach solution (made using Cleanline thin bleach containing 4.53% sodium hypochlorite), followed by 70% v/v ethanol solution. Natural samplers, eDNA filters, DNA extracts and PCR reagents were each stored in separate freezers. All PCR preparation was conducted inside a hood sterilised with ultraviolet (UV) light and 10% v/v bleach solution. All plastics were purchased sterile but additionally sterilised inside the hood with UV light for at least 30 minutes. Metal forceps (Fisher Scientific, UK) and dissection scissors (Fisher Scientific, UK) were sterilised in a 10% v/v bleach bath for 5 minutes, then immersed in a 5% v/v Lipsol detergent (Scientific Laboratory Supplies, UK) bath and rinsed with deionised water. These were further sterilised inside the hood with UV light for at least 30 minutes before use.

Stainless steel 5 mm beads (Qiagen, Germany) were sterilised by rinsing with deionised water, immersion in 10% bleach v/v solution in a 50 ml falcon tube for 1 minute, then immersion in 70% v/v ethanol solution in a 50 ml falcon tube for 1 minute. The beads were transferred to a 500 ml glass conical flask and the flask opening was covered with aluminium foil. The flask was baked at 220°C for 3 hours and allowed to cool before beads were transferred to a new 50 ml falcon tube.

All other reusable equipment (e.g. tube racks, glassware) was sterilised in a 10% v/v bleach bath for at least 10 minutes, then immersed in a 5% v/v Lipsol detergent bath and rinsed with deionised water. Bleach and Lipsol detergent baths were changed weekly or daily during periods of heavy use. The entire laboratory was deep-cleaned (benches and equipment sterilised, waste discarded, floors swept and mopped with 10% v/v bleach solution) on a weekly basis.

Prepared PCR reactions were transported to a separate laboratory for addition of PCR positive controls and PCR amplification. PCR products were transported to another laboratory for gel electrophoresis, library preparation, and storage. Library quality checks, quantification, and sequencing were performed in a separate laboratory. Bench space and equipment in all laboratories were sterilised with 10% v/v bleach solution, followed by 70% v/v ethanol solution.

#### Database creation

*Aquarium species*

All available mitochondrial 12S records for stocked fish species and the PCR positive control (*Pangasius hypophthalmus*) were downloaded from GenBank (Table S1).

*Marine vertebrates*

For increased speed and accuracy during taxonomic assignment, rather than using the entirety of the GenBank nucleotide database, we created a reduced database of marine vertebrates. All available mitochondrial 12S records for specific marine vertebrate taxonomic ranks were downloaded from GenBank: seals/sea otters/otters/polar bears (Odobenidae, Otariidae, Phocidae, *Enhydra lutris*, *Lutra lutra*, *Ursus maritimus*), manatees (Sirenia), whales and dolphins (Cetacea), fish (Actinopterygii, Agnatha, Elasmobranchii, Holocephali, Latimeria, Petromyzontiformes), birds (Aves) and reptiles (Hydrophiinae, Testudines, *Crocodylus porosus*).

*Domestic species*

Both reference libraries also included the first 10 GenBank records of common domestic species (cow; *Bos taurus*, pig; *Sus scrofa*, chicken; *Gallus gallus*, sheep; *Ovis aries*, dog; *Canis lupus*, and cat; *Felis catus*). Additionally, we included a specific set of human (*Homo sapiens*) 12S records.

*Final database creation*

In total, 52,804 records for 14,382 unique species were used to create the final marine vertebrate BLAST database, and 183 records for 34 unique species were used to create the final aquarium BLAST database.

### Results

#### Phase 1

*Total DNA yield*
For *L. chondrodes*, extraction protocol combined with ethanol treatment ($F_8$ = 0.220, P = 0.985) did not influence total DNA yield. Without an interaction term, ethanol treatment ($F_2$ = 13.427, P < 0.001) and extraction protocol ($F_4$ = 6.080, P = 0.001) both influenced total DNA yield. The model with no interaction term was not overdispersed ($\theta_{38}$ = 0.873, P = 0.692) but the residuals deviated from normality (Shapiro-Wilk normality test: W = 0.875, P < 0.001). The data were split by ethanol treatment, and Kruskal-Wallis Test and Dunn's Test used to examine the effect of extraction protocol. Total DNA yield significantly differed between extraction protocols for the spin ethanol treatment ($H_4$ = 11.314, P = 0.023), but not the blot ($H_4$ = 8.833, P = 0.065) or wet ($H_4$ = 6.754, P = 0.150) treatments. For the spin treatment, none of the Dunn's Test adjusted p-values were significant, but the largest differences in effect size were observed between the QBT and MUS protocols (Z = -2.749, P = 0.060), the QBT and MUT protocols (Z = -2.107, P = 0.117), and the QBT and QPS protocols (Z = 2.565, P = 0.052). The data were also split by extraction protocol to compare ethanol treatments using Kruskal-Wallis Test and Dunn's Test. Total DNA yield significantly differed between ethanol treatments with the QBT ($H_2$ = 7.200, P = 0.027) and MUT ($H_2$ = 7.261, P = 0.027) protocols. For both protocols, the spin treatment produced lower total DNA yields than the wet treatment (QBT: Z = -2.683, P = 0.022; MUT: Z = -2.695, P = 0.021), but the blot treatment did not significantly differ to the spin or wet treatments. No differences between ethanol treatments were found with other extraction protocols (MUS: $H_2$ = 5.600, P = 0.061; QBT-ZYMO: $H_2$ = 0.964, P = 0.618; QPS: $H_2$ = 3.822, P = 0.148).

For *V. pourtalesii*, extraction protocol combined with ethanol treatment ($F_8$ = 1.194, P = 0.335) did not influence total DNA yield. Without an interaction term, ethanol treatment did not influence total DNA yield ($F_2$ = 2.289, P = 0.115) but extraction protocol did ($F_4$ = 29.312, P < 0.001). The model with no interaction term was not overdispersed ($\theta_{38}$ = 0.921, P = 0.609) but the residuals deviated from normality (Shapiro-Wilk normality test: W = 0.865, P < 0.001). The data were split by ethanol treatment and Kruskal-Wallis Test with Dunn's Test used to examine the effect of extraction protocol. Total DNA yield significantly differed between extraction protocols for the wet ($H_4$ = 13.233, P = 0.010) and blot ($H_4$ = 11.833, P = 0.019) ethanol treatments, but not the spin ($H_4$ = 8.233, P = 0.083) treatment. The QBT protocol produced higher total DNA yields than the QPS protocol with the wet ethanol treatment (Z = 3.286, P = 0.010) and the MUT protocol with the blot treatment (Z = -2.921, P = 0.035).

*Post-PCR DNA concentration*
For *L chondrodes*, extraction protocol combined with ethanol treatment ($F_8$ = 1.425, P = 0.227) did not influence post-PCR DNA concentration. Without an interaction term, extraction protocol ($F_4$ = 2.350, P = 0.072) did not influence post-PCR DNA concentration, but ethanol treatment did ($F_2$ = 24.188, P < 0.001). The model with no

interaction term was overdispersed ($\theta_{38}$ = 4.827, P < 0.001) and the residuals deviated from normality (Shapiro-Wilk normality test: W = 0.950, P = 0.033). The data were split by extraction protocol, and Kruskal-Wallis Test and Dunn's Test used to examine the effect of ethanol treatment. Ethanol treatment was found to influence post-PCR DNA concentration with the MUS protocol ($H_2$ = 6.006, P = 0.050), but not other protocols (MUT: $H_2$ = 5.067, P = 0.079; QBT: $H_2$ = 5.422, P = 0.067; QBT-ZYMO: $H_2$ = 1.501, P = 0.472; QPS: $H_2$ = 5.600, P = 0.061). For the MUS protocol, the spin treatment produced lower post-PCR DNA concentrations than the wet treatment (Z = -2.395, P = 0.050), but the blot treatment did not significantly differ to the spin or wet treatments.

For *V. pourtalesii*, extraction protocol combined with ethanol treatment ($F_8$ = 2.012, P = 0.079) did not influence post-PCR DNA concentration. Without an interaction term, ethanol treatment did not influence post-PCR DNA concentration ($F_2$ = 0.641, P = 0.533), but extraction protocol did ($F_4$ = 2.964, P = 0.032). The model with no interaction term was overdispersed ($\theta_{38}$ = 4.314, P < 0.001) but all other assumptions were met. Post-PCR DNA concentration was positively influenced by the MUT (0.669 ± 0.327, t = 2.044, P = 0.048) and QBT (1.005 ± 0.311, t = 3.229, P = 0.003) protocols, but the MUT protocol (12.13 ± 2.31 ng/µl) produced lower post-PCR DNA concentrations than the QBT protocol (16.97 2.74 ng/µl; Z = -3.229, P = 0.011). No differences were found between other extraction protocols.

*Taxon richness*

For *L. chondrodes*, extraction protocol combined with ethanol treatment ($\chi_8$ = 8.416, P = 0.394) did not influence fish taxon richness. Without an interaction term, extraction protocol ($\chi_4$ = 2.076, P = 0.722) and ethanol treatment ($\chi_2$ = 1.121, P = 0.571) did not influence fish taxon richness. The model with no interaction term was not found to be overdispersed ($\theta_{30}$ = 0.355, P = 0.999) but the residuals deviated from normality (Shapiro-Wilk normality test: W = 0.946, P = 0.035).

For *V. pourtalesii*, extraction protocol combined with ethanol treatment ($\chi_8$ = 13.863, P = 0.085) did not influence fish taxon richness. Without an interaction term, extraction protocol ($\chi_4$ = 12.240, P = 0.016) influenced fish taxon richness, but ethanol treatment ($\chi_2$ = 2.758, P = 0.252) did not. The model with no interaction term was not found to be overdispersed ($\theta_{38}$ = 1.025, P = 0.427) but the residuals deviated from normality (Shapiro-Wilk normality test: W = 0.743, P < 0.001). The data were split by ethanol treatment, and Kruskal-Wallis Test and Dunn's Test used to examine the effect of extraction protocol. Extraction protocol was not found to influence fish taxon richness for the spin ($H_4$ = 5.343, P = 0.254), blot ($H_4$ = 6.990, P = 0.136) or wet ($H_4$ = 3.238, P = 0.519) ethanol treatments.

*Proportional read counts*

For *L. chondrodes*, extraction protocol combined with ethanol treatment ($F_8$ = 2.299, P = 0.047) influenced fish proportional read counts. Without an interaction term, extraction protocol ($F_4$ = 2.488, P = 0.060) did not influence fish proportional read counts, but ethanol treatment ($F_2$ = 15.476, P < 0.001) did. The model with no interaction term was not found to be overdispersed ($\theta_{38}$ = 0.533, P = 0.992) but the residuals deviated from normality (Shapiro-Wilk normality test: W = 0.887, P < 0.001). The data were split by extraction protocol, and Kruskal-Wallis Test and Dunn's Test used to examine the effect of ethanol treatment. Ethanol treatment was not found to influence proportional read counts for the MUT ($H_2$ = 5.956, P = 0.051), QBT ($H_2$ = 5.422, P = 0.067) or QBT-ZYMO ($H_2$ = 4.146, P = 0.126) protocols. Proportional read counts significantly differed between ethanol treatments with the MUS ($H_2$ = 6.161, P = 0.046) and QPS ($H_2$ = 6.006, P = 0.050) protocols. The wet treatment produced higher proportional read counts than the spin treatment with the MUS protocol (Z = -2.426, P = 0.046), and than the blot treatment with the QPS protocol (Z = -2.395, P = 0.050). No significant differences were found between other treatments with either protocol or between ethanol treatments with other extraction protocols (MUT: $H_2$ = 5.956, P = 0.051; QBT: $H_2$ = 5.422, P = 0.067; QBT-ZYMO: $H_2$ = 4.146, P = 0.126).

For *V. pourtalesi*, extraction protocol combined with ethanol treatment ($F_8$ = 1.696, P = 0.141) did not influence fish proportional read counts. Without an interaction term, both extraction protocol ($F_4$ = 3.671, P = 0.013) and ethanol treatment ($F_2$ = 8.446, P < 0.001) influenced fish proportional read counts. The model with no interaction term was not found to be overdispersed ($\theta_{38}$ = 0.317, P = 0.999) but the residuals deviated from normality (Shapiro-Wilk normality test: W = 0.866, P < 0.001). The data were split by ethanol treatment, and Kruskal-Wallis Test and Dunn's Test used to examine the effect of extraction protocol. Extraction protocol was not found to influence proportional read counts for the spin ($H_4$ = 5.326, P = 0.256), blot ($H_4$ = 7.598, P = 0.108) or wet ($H_4$ = 3.238, P = 0.519) ethanol treatments. The data were also split by extraction protocol to compare ethanol treatments using Kruskal-Wallis Test and Dunn's Test. Proportional reads counts significantly differed between ethanol treatments with the QBT-ZYMO protocol ($H_2$ = 7.624, P = 0.022), but not other DNA extraction protocols (MUS: $H_2$ = 1.167, P = 0.558; MUT: $H_2$ = 1.818, P = 0.403; QBT: $H_2$ = 1.167, P = 0.558; QPS: $H_2$ = 2.000, P = 0.368). The blot treatment produced higher proportional read counts than the spin (Z = 2.391, P = 0.025) or wet (Z = 2.391, P = 0.050) treatments. No significant differences were found between the spin and wet treatments.

#### Phase 2

*Total DNA yield*

For *P. ventilabrum*, type of starting material combined with amount ($F_2$ = 3.206, P = 0.053) did not influence total DNA yield. Without an interaction term, type ($F_1$ = 14.641, P < 0.001) and amount ($F_2$ = 16.485, P < 0.001) of starting material both influenced total DNA yield. The model with no interaction term was overdispersed ($\theta_{37}$ = 5.055, P < 0.001) but all other assumptions were met. Total DNA yield was higher using homogenised tissue (0.460 ± 0.120, t = 3.840, P < 0.001) and increased as the amount of starting material increased (amount 2: 0.451 ± 0.136, t = 3.321, P = 0.002; amount 3: 0.981 ± 0.165, t = 5.931, P < 0.001). Dried tissue (30.30 ± 2.81 ng/µl) produced lower total DNA yields than homogenate (48.00 ± 3.82 ng/µl; Z = -3.840, P < 0.001). Amount 3 (63.10 ± 7.96 ng/µl) produced higher total DNA yields than amount 2 (37.10 ± 3.18 ng/µl; Z = -3.474, P = 0.002) or amount 1 (23.70 ± 2.53 ng/µl; Z = -5.931, P < 0.001). Amount 1 also produced lower total DNA yields than amount 2 (Z = -3.321, P = 0.003).

For *V. pourtalesii*, type of starting material combined with amount ($F_2$ = 0.676, P = 0.514) did not influence total DNA yield. Without an interaction term, type of starting material ($F_1$ = 0.433, P = 0.514) did not influence total DNA yield, but amount ($F_2$ = 3.795, P = 0.030) did. The model with no interaction term was overdispersed ($\theta_{48}$ = 0.488, P < 0.001) and the residuals deviated from normality (Shapiro-Wilk normality test: W = 0.928, P = 0.004). The data were split by type of starting material, and Kruskal-Wallis Test and Dunn's Test used to examine the effect of amount of starting material. Amount of starting material was not found to influence total DNA yield for homogenised ($H_2$ = 4.936, P = 0.085) or dried ($H_2$ = 2.181, P = 0.336) tissue.

*Post-PCR DNA concentration*

For *P. ventilabrum*, type of starting material combined with amount ($F_2$ = 0.420, P = 0.660) did not influence post-PCR DNA concentration. Without an interaction term, type of starting material ($F_1$ = 5.054, P = 0.031) influenced post-PCR DNA concentration, but amount did not ($F_2$ = 0.697, P = 0.504). The model with no interaction term was overdispersed ($\theta_{37}$ = 2.003, P < 0.001) and the residuals deviated from normality (Shapiro-Wilk normality test: W = 0.916, P = 0.005). The data were split by amount of starting material, and Kruskal-Wallis Test and Dunn's Test used to examine the effect of type of starting material. Type of starting material was not found to influence post-PCR concentration for amount 1 ($H_1$ = 0.860, P = 0.354), amount 2 ($H_1$ = 3.608, P = 0.058) or amount 3 ($H_1$ = 1.333, P = 0.248).

For *V. pourtalesii*, type of starting material combined with amount ($F_2$ = 0.535, P = 0.589) did not influence post-PCR DNA concentration. Without an interaction term, type ($F_1$ = 0.652, P = 0.423) and amount ($F_2$ = 2.712, P = 0.077) of starting material did not influence post-PCR concentration. The model with no interaction term was overdispersed ($\theta_{48}$ = 5.806, P < 0.001) and the residuals deviated from normality (Shapiro-Wilk normality test: W = 0.950, P = 0.028).

*Taxon richness*

For *P. ventilabrum*, type of starting material combined with amount ($\chi_2$ = 0.025, P = 0.988) did not influence fish taxon richness. Without an interaction term, type ($\chi_1$ = 0.111, P = 0.739) and amount ($\chi_2$ = 0.590, P = 0.744) of starting material did not influence fish taxon richness. The model without the interaction term was not overdispersed ($\theta_{37}$ = 0.106, P = 1.000) and all assumptions were met.

For *V. pourtalesii*, type of starting material combined with amount ($\chi_2$ = 0.439, P = 0.803) did not influence fish taxon richness. Without an interaction term, type of starting material ($\chi_1$ = 6.451, P = 0.011) influenced fish taxon richness, but amount ($\chi_2$ = 2.168, P = 0.338) did not. The model without the interaction term was not overdispersed ($\theta_{48}$ = 1.163, P = 0.204) and all assumptions were met. Type of starting material had a positive influence on taxon richness (0.476 ± 0.190, t = 2.514, P = 0.012), where taxon richness was higher using homogenate (2.780 ± 0.334) than dried tissue (1.720 ± 0.253; Z = -2.514, P = 0.012).

*Proportional read counts*

For *P. ventilabrum*, type of starting material combined with amount ($F_2$ = 1.164, P = 0.324) did not influence fish proportional read counts. Without an interaction term, type ($F_1$ = 0.734, P = 0.397) and amount ($F_2$ = 2.495, P = 0.096) of starting material did not influence fish proportional read counts. The model without an interaction term was not overdispersed ($\theta_{37}$ = 0.030, P = 0.980) and all assumptions were met.

For *V. pourtalesii*, type of starting material combined with amount ($F_2$ = 0.103, P = 0.902) did not influence fish proportional read counts. Without an interaction term, type of starting material ($F_1$ = 3.966, P = 0.052) did not influence fish proportional read counts, but amount ($F_2$ = 3.266, P = 0.047) did. The model without an interaction term was not overdispersed ($\theta_{48}$ = 0.062, P = 1.000) but the residuals deviated from normality (Shapiro-Wilk normality test: W = 0.866, P < 0.001). The data were split by type of starting material, and Kruskal-Wallis Test and Dunn's Test used to examine the effect of amount of starting material. Amount of starting material was not found to influence proportional read counts for homogenate ($H_2$ = 3.705, P = 0.157) or dried tissue ($H_2$ = 1.147, P = 0.564).

### Tables

**Table S1.** Marine aquarium species list. Species marked with '*' had no 12S records available on GenBank.

| **Aquarium species present** |
|---|
| *Acanthurus guttatus* |
| *Amphiprion frenatus* |
| *Amphiprion ocellaris* |
| *Amphiprion percula* |
| *Centropyge loricula* |
| *Chelmon rostratus* |
| *Chromis vanderbilti* |
| *Chromis viridis* |
| *Chromis xanthura* |
| *Chrysiptera parasema* |
| *Forcipiger flavissimus* |
| *Genicanthus bellus* |
| *Gobiodon okinawae* |
| *Gramma loreto* |
| *Halichoeres melasmapomus* |
| *Hippocampus semispinosis* * |
| *Naso elegans* |
| *Oxycirrhites typus* |
| *Pangasius hypophthalmus* |
| *Paracanthurus hepatus* |
| *Pictichromis porphyrea* |
| *Pomacanthus imperator* |
| *Pseudanthias cheirospilos* * |
| *Pseudanthias pleurotaenia* |
| *Pseudocheilinus hexataenia* |
| *Ptereleotris zebra* |
| *Scarus quoyi* |
| *Siganus vulpinus* |
| *Zebrasoma flavescens* |
| *Zebrasoma veliferum* |

### Figures

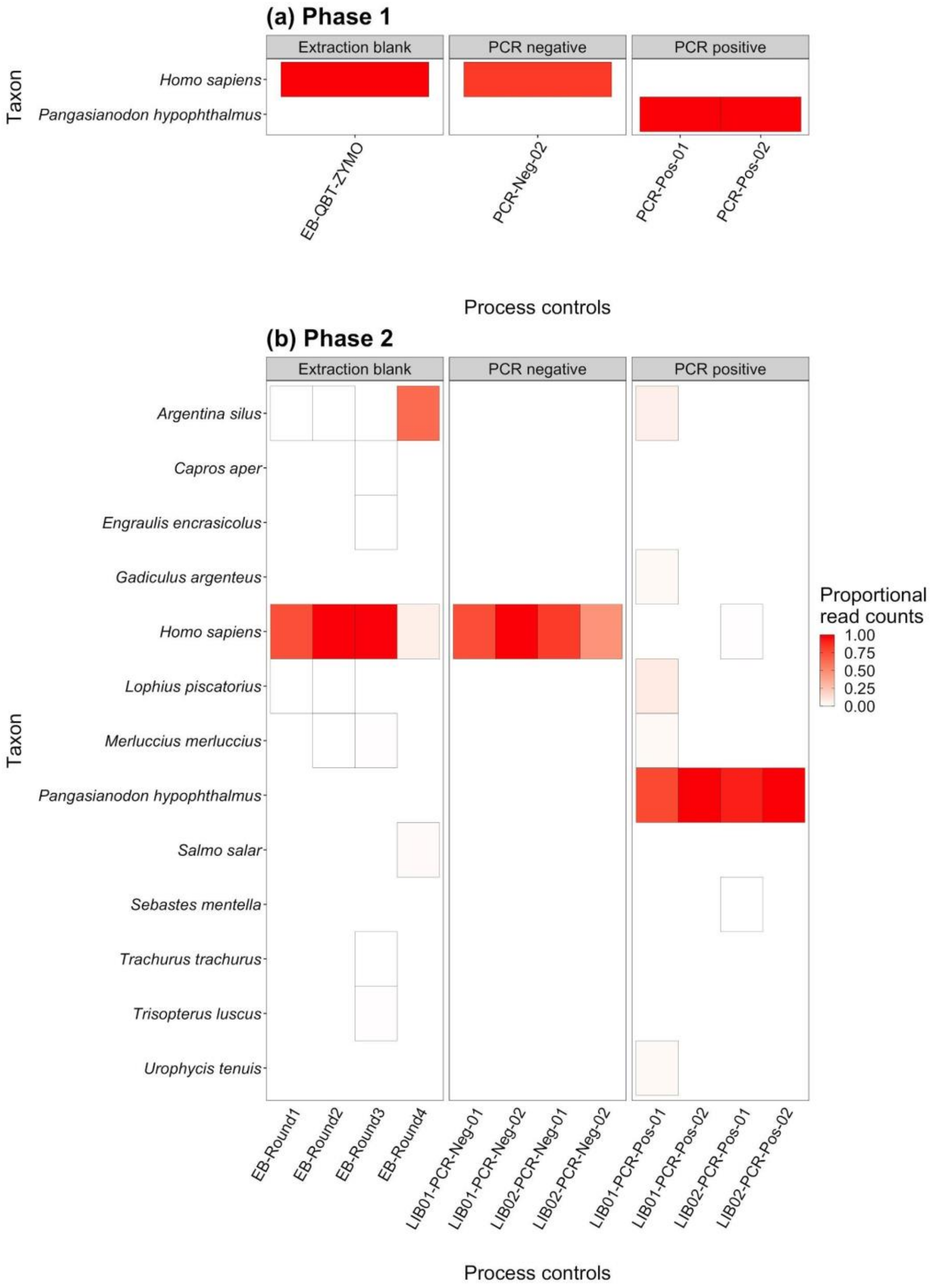


**Figure S1.** Heatmap showing the frequency of contaminants detected in process controls with sequence reads. Assignments that were not detected in a contaminated control are coloured white with no border.

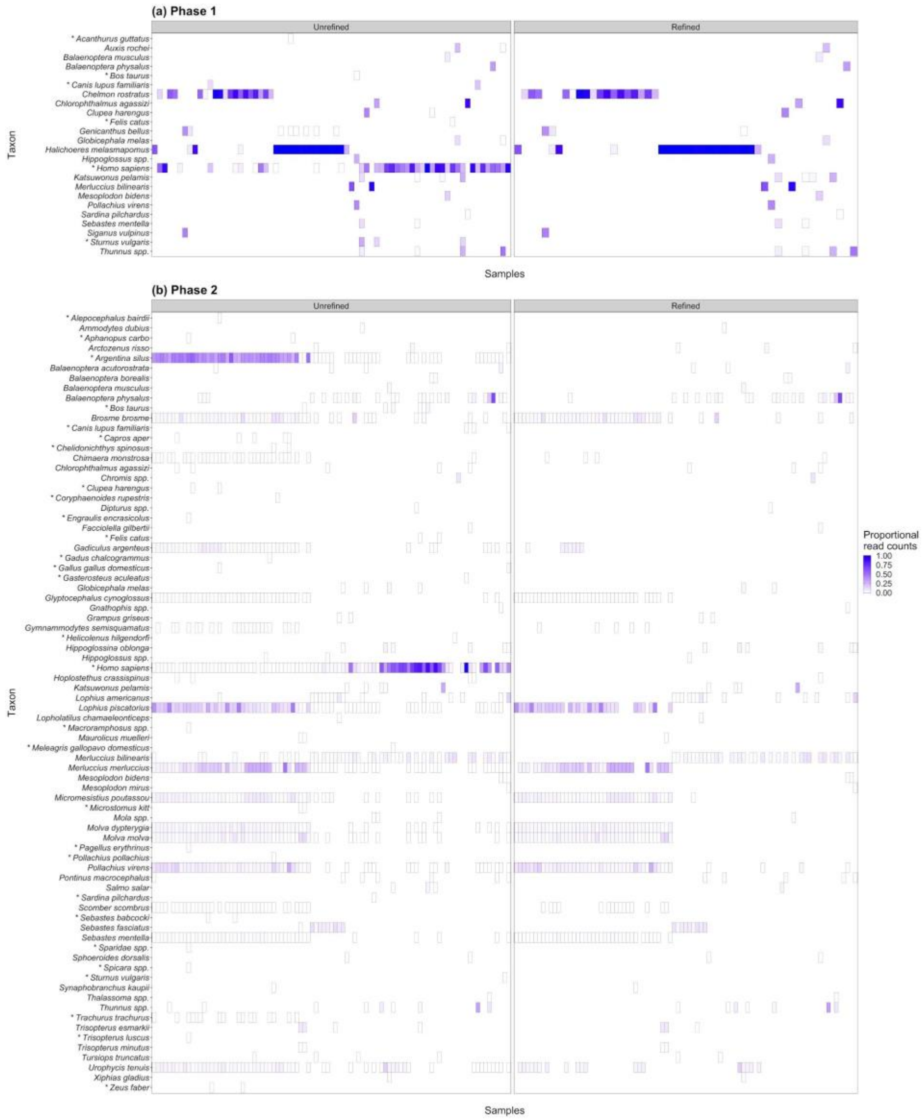


**Figure S2.** Heatmap showing proportional read counts for taxa detected in experimental Phase 1 **(a)** and Phase 2 **(b)** sponge samples before and after dataset refinement, which included application of taxon-specific sequence thresholds to remove false positives arising through contamination (Phase 2 data only) and a low-noise sequence threshold (0.01%) to remove false positives arising from taxa occurring at low frequency, as well as removal of higher taxonomic ranks, iridescent shark (*Pangasionodon hypopthalmus* - PCR positive control), human, domestic species (*Bos taurus*, *Canis lupus familiaris*, *Felis catus*, *Gallus gallus domesticus*, *Meleagris gallopavo domesticus*) and terrestrial species (*Sturnus vulgaris*). Symbols after taxon names indicate taxa that were removed during dataset refinement. Taxon names with no symbol indicate taxa retained by dataset refinement for downstream analyses. Taxa that were not detected in a sample before or after dataset refinement are coloured white with no border.

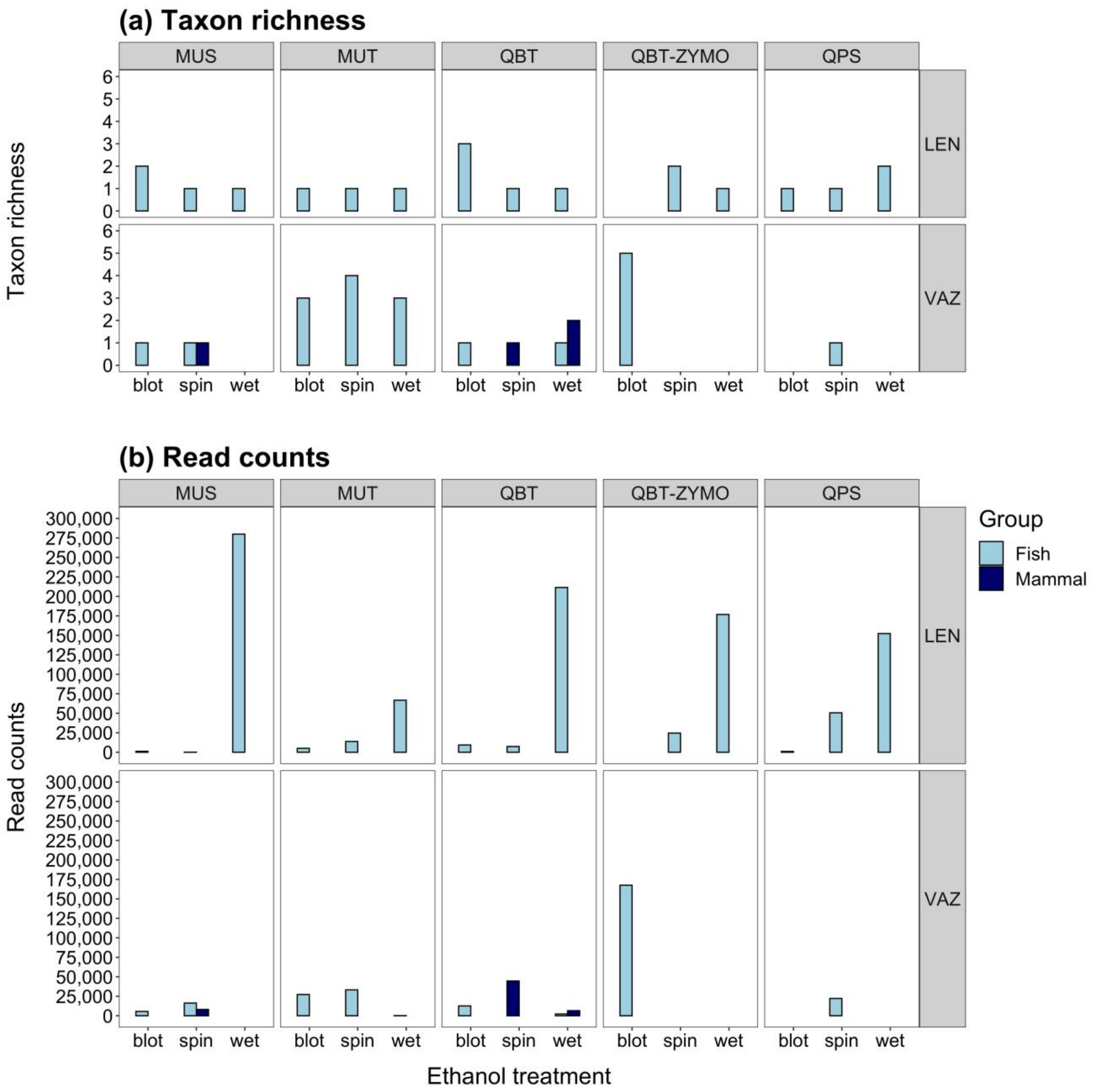


**Figure S3.** Bar plot showing fish and mammal **(a)** taxon richness and **(b)** read counts for samples processed with each ethanol treatment (blot, spin, wet) and DNA extraction protocol (MUS, MUT, QBT, QBT-ZYMO and QPS) for different sponge species in experimental Phase 1. Abbreviations: *Lendenfeldia chondrodes* (LEN), *Vazella pourtalesii* (VAZ).

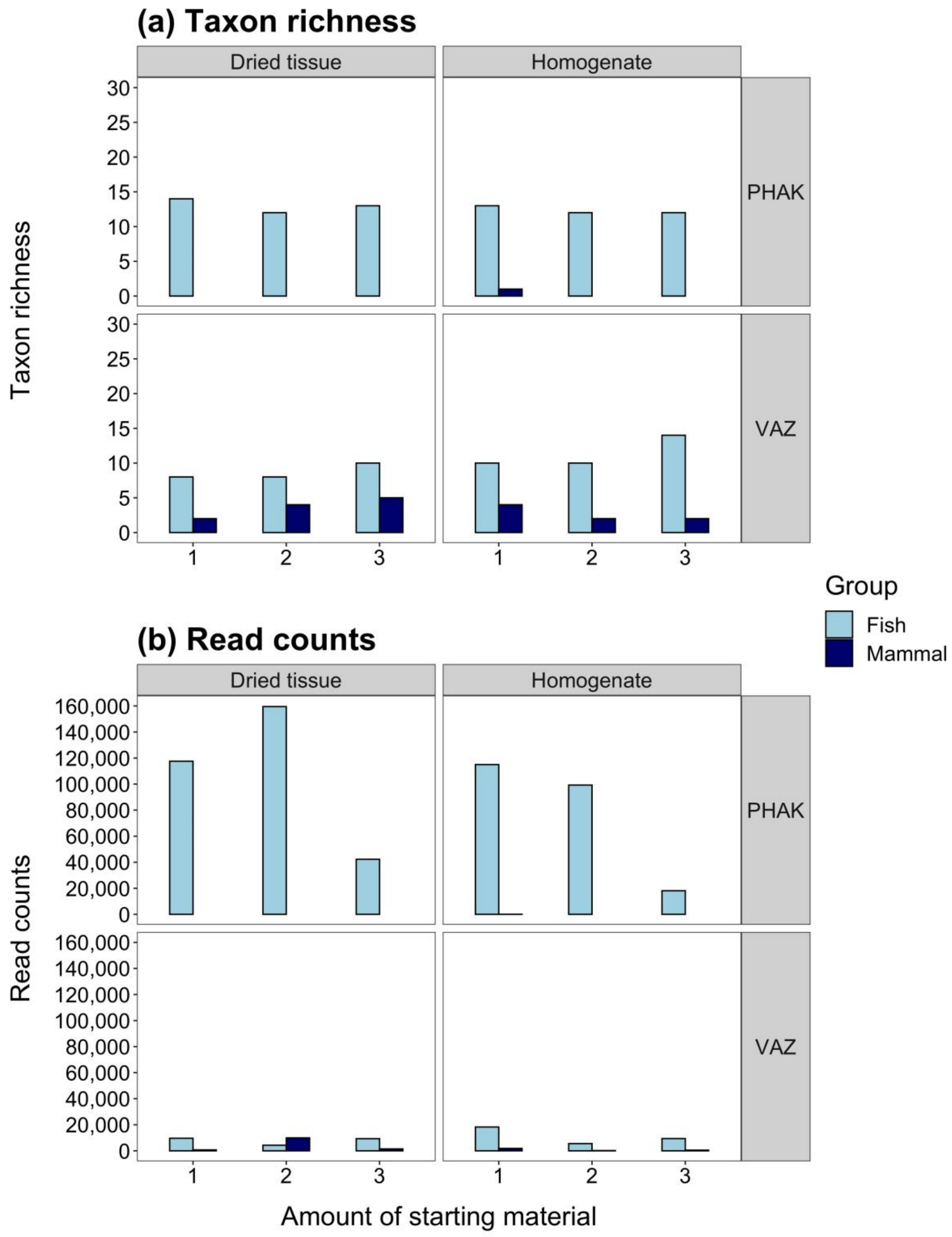


**Figure S4.** Bar plot showing fish and mammal **(a)** taxon richness and **(b)** read counts for samples of each type and amount of starting material for different sponge species in experimental Phase 2. Abbreviations: *Phakellia ventilabrum* (PHAK), *Vazella pourtalesii* (VAZ).

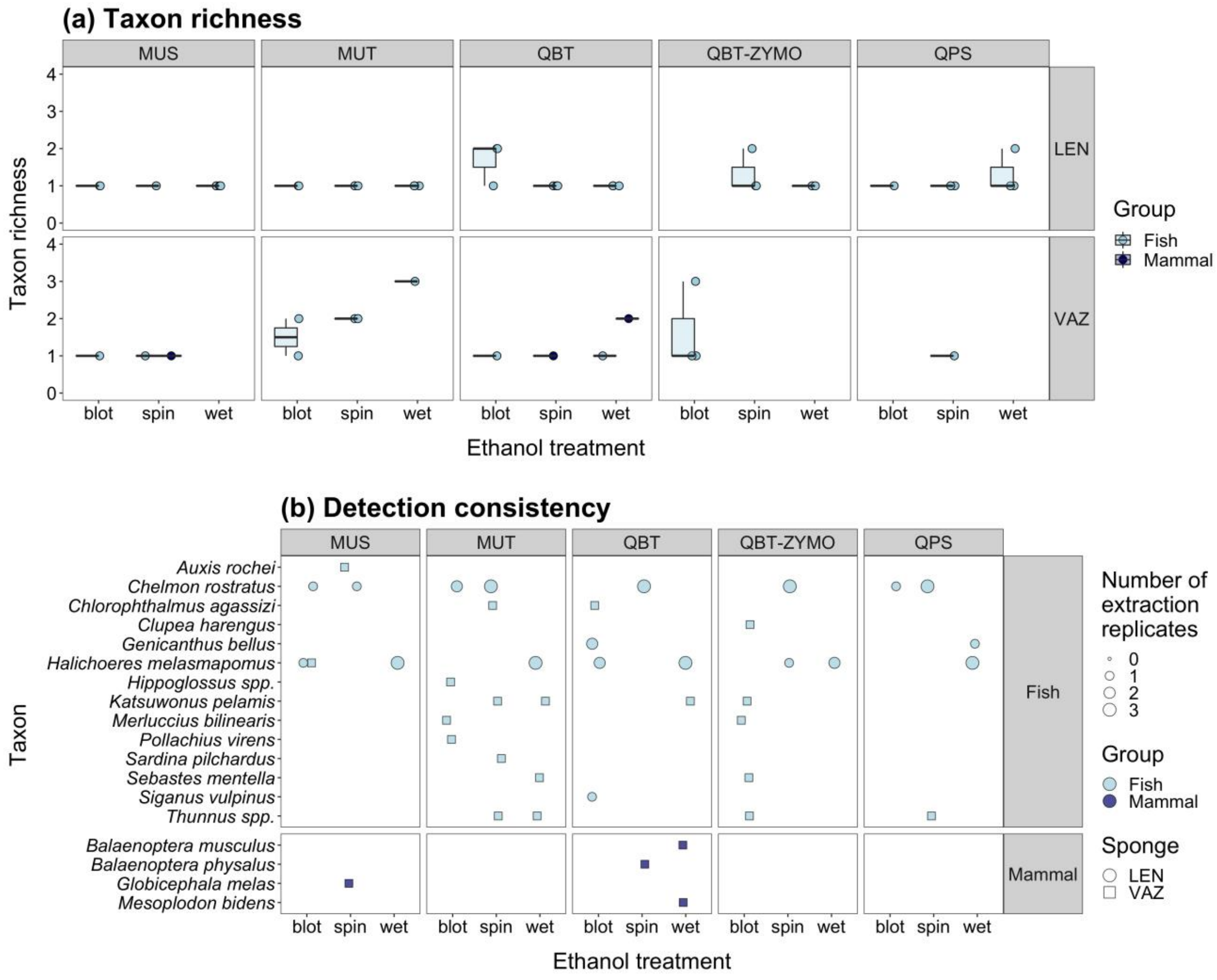


**Figure S5.** Summary of detection consistency across extraction replicates for sponge samples in experimental Phase 1. The box plot shows taxon richness of each extraction replicate for samples processed with each ethanol treatment (blot, spin, wet) and DNA extraction protocol (MUS, MUT, QBT, QBT-ZYMO and QPS) **(a)**, and the bubble plot shows the number of extraction replicates for each ethanol treatment and DNA extraction protocol where a given taxon was detected **(b)**. Abbreviations: *Lendenfeldia chondrodes* (LEN), *Vazella pourtalesii* (VAZ).

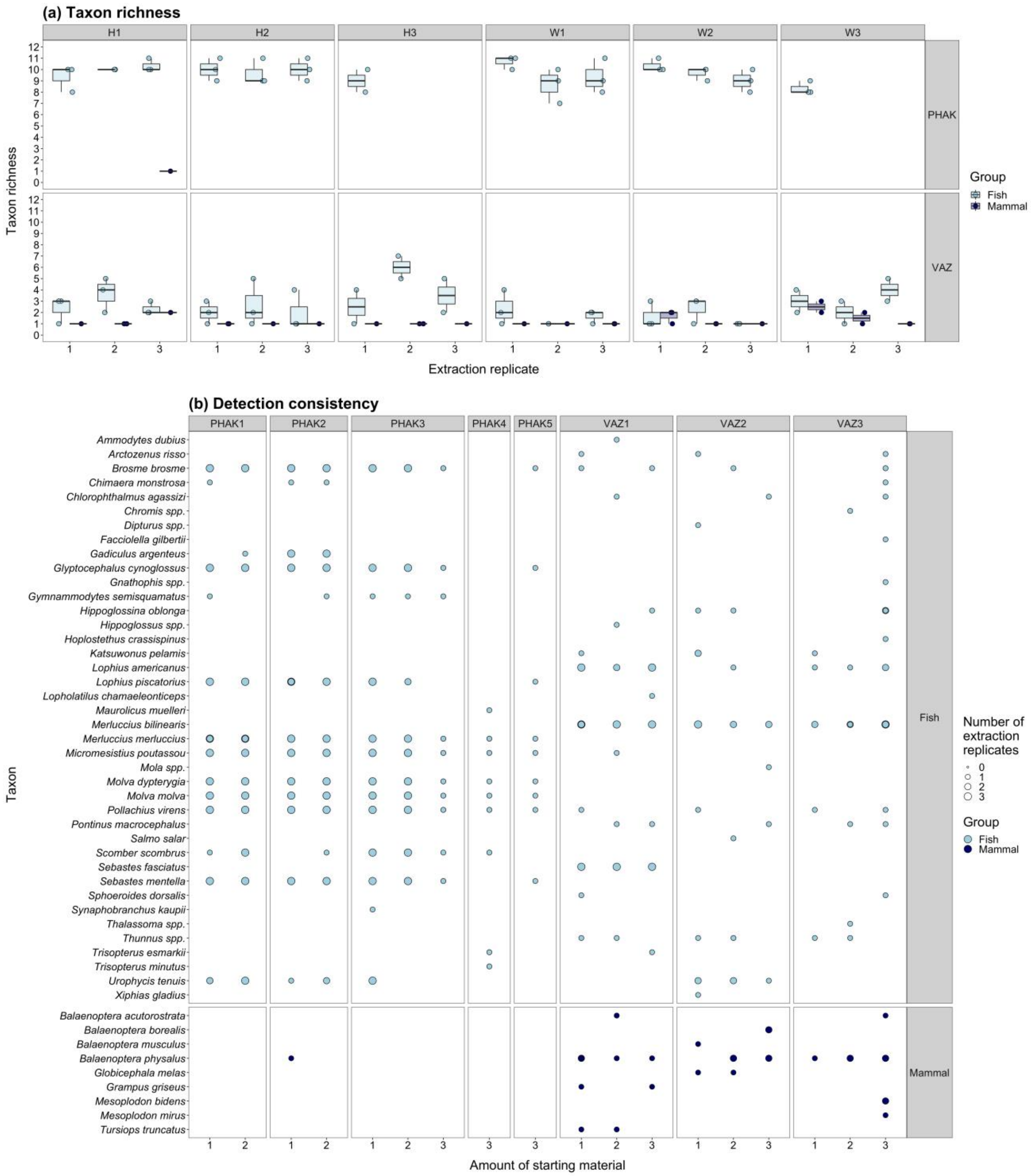


**Figure S6.** Summary of detection consistency across extraction replicates for *Vazella pourtalesii* and *Phakellia ventilabrum* samples in experimental Phase 2. The box plot shows taxon richness of each extraction replicate for each amount of starting material **(a)**, and the bubble plot shows the number of extraction replicates for each type and amount of starting material where a given taxon was detected **(b)**. Abbreviations: *Phakellia ventilabrum* (PHAK), *Vazella pourtalesii* (VAZ).
